# Distinct progenitor behavior underlying neocortical gliogenesis related to tumorigenesis

**DOI:** 10.1101/2020.10.13.338459

**Authors:** Zhongfu Shen, Yang Lin, Jiajun Yang, David J. Jörg, Yuwei Peng, Xiuli Zhang, Yifan Xu, Luisirene Hernandez, Jian Ma, Benjamin D. Simons, Song-Hai Shi

## Abstract

Radial glial progenitors (RGPs) are responsible for producing the vast majority of neurons and glia in the neocortex. While RGP behavior and progressive generation of neocortical neurons have been delineated, the exact process of neocortical gliogenesis remains elusive. Here, we report the precise progenitor cell behavior and gliogenesis program at single-cell resolution in the mouse neocortex. RGPs transition from neurogenesis to gliogenesis progressively, producing astrocytes, oligodendrocytes, or both in well-defined propensities of 60%:15%:25%, respectively, via fate-restricted “intermediate” precursor cells. While the total number of precursor cells generated by individual RGPs appears stochastic, the output of individual precursor cells exhibit clear patterns in number and subtype, and form discrete local subclusters. Clonal loss of tumor suppressor Neurofibromatosis type 1 leads to excessive production of glia selectively, especially oligodendrocyte precursor cells. These results delineate the cellular program of neocortical gliogenesis quantitatively and suggest the cellular and lineage origin of primary brain tumor.

## INTRODUCTION

The neocortex is a hallmark of mammals and the seat of higher-order brain functions, such as sensory perception, movement, language and cognition. It consists of a large number of neurons and glial cells that are organized into distinct laminae. Previous studies have shown that radial glia progenitors (RGPs) are responsible for producing nearly all neurons and glia in the neocortex (Anthony et al., 2004; Costa et al., 2009; Florio and Huttner, 2014; Gao et al., 2014; Homem et al., 2015; Kriegstein and Alvarez-Buylla, 2009; Malatesta et al., 2000; Miyata et al., 2001; Noctor et al., 2001; Noctor et al., 2004; Rowitch and Kriegstein, 2010; Tamamaki et al., 2001). During the early stage of neocortical development, proliferation of neuroepithelial cells leads to the generation of radial glial cells, the predominant neural progenitor cells in the developing neocortex. RGPs reside in the ventricular zone and possess a characteristic bipolar morphology (Bultje et al., 2009; Chenn et al., 1998; Rakic, 2003). Notably, RGPs conform to a highly regulated and deterministic program transitioning sequentially through distinct phases (Gao et al., 2014). Initially, RGPs move through an amplification phase, expanding their number through a series of symmetric divisions. Subsequently, RGPs transition into a neurogenic phase where they undergo predominantly asymmetric division giving rise to neurons either directly or indirectly via transit amplifying progenitors, such as intermediate progenitor cells (Englund et al., 2005; Haubensak et al., 2004; Miyata et al., 2004) or outer subventricular zone RGPs (also called basal or intermediate RGPs; more abundant in primates) (Betizeau et al., 2013; Fietz et al., 2010; Hansen et al., 2010; Reillo et al., 2011; Shitamukai et al., 2011; Wang et al., 2011). Newborn neurons migrate radially to constitute the future neocortex in a birth date-dependent inside-out manner (Angevine and Sidman, 1961; Hatten, 1999; Marin and Rubenstein, 2003; Noctor et al., 2001; Rakic, 1988). Proper RGP behavior, and orderly neurogenesis and neuronal migration are fundamental to neocortical development and laminar formation.

Alongside neurons, glial cells are vital components of the neocortex and play crucial roles in supporting neocortical development and function, including blood brain barrier formation and maintenance, synaptogenesis, neurotransmission, metabolic regulation, as well as modulation and support of synaptic transmission (Allen and Barres, 2009; Barres, 2008; Bergles and Richardson, 2015; Freeman and Rowitch, 2013; Molofsky et al., 2012; Rowitch and Kriegstein, 2010; Stogsdill and Eroglu, 2017). There are two major classes of glial cells in the neocortex, macroglia and microglia (Reemst et al., 2016). Microglia are the resident immune cells and stem from the mesoderm (Alliot et al., 1999), colonizing the developing neocortex during early embryonic development (Swinnen et al., 2013). Macroglia including astrocytes and oligodendrocytes share the same progenitor origin as neurons (Kessaris et al., 2006; Kriegstein and Alvarez-Buylla, 2009; Noctor et al., 2004; Rowitch and Kriegstein, 2010; Tabata, 2015). Previous studies suggest that a subset of RGPs proceed to gliogenesis to produce astrocytes and oligodendrocytes in the neocortex (Clavreul et al., 2019; Gao et al., 2014; Garcia-Marques and Lopez-Mascaraque, 2013; Magavi et al., 2012; Siddiqi et al., 2014). However, the precise behavior of RGPs in transitioning to gliogenesis and the cellular program underlying neocortical gliogenesis remain largely unknown.

Glial cells in the neocortex are diverse, as reflected in their morphology, location, molecular expression, biophysical properties and function (Bayraktar et al., 2014; Bergles and Richardson, 2015; Emery and Dugas, 2013; Emsley and Macklis, 2006; Garcia-Marques and Lopez-Mascaraque, 2013; John Lin et al., 2017; Lanjakornsiripan et al., 2018; Lee et al., 2006; Marques et al., 2016; Middeldorp and Hol, 2011; Molofsky and Deneen, 2015; Schitine et al., 2015; Silbereis et al., 2010; Suzuki et al., 2017; Tabata, 2015; Zeisel et al., 2018; Zhang et al., 2016). For example, layer 1 astrocytes (L1A, also referred to as the glia limitans underneath the pia astrocytes) are preferentially located in layer 1 with a flat cell body and short processes, and exhibit distinct spontaneous Ca^2+^ signaling activity and molecular expression (Garcia-Marques and Lopez-Mascaraque, 2013; Liu et al., 2013; Takata and Hirase, 2008; Zeisel et al., 2018). The vast majority of astrocytes in layers II-VI are often collectively referred to as protoplasmic astrocytes (PA), even though recent studies have indicated the existence of molecularly distinct subpopulations (Bayraktar et al., 2020; Lanjakornsiripan et al., 2018; Zeisel et al., 2018). They possess short and highly branched tertiary processes. In addition, the fibrous astrocytes (FA) are usually located within the white matter and grow relatively long unbranched processes. Oligodendrocytes arise from oligodendrocyte precursor cells (OPCs), which can be identified by their expression of a number of proteins, such as the NG2 chondroitin sulfate proteoglycan and the platelet-derived growth factor α receptor (PDGFRα) (Emery and Dugas, 2013; Suzuki et al., 2017). After birth, oligodendrocytes progressively mature into myelin-forming oligodendrocytes, while many OPCs persist in the adult neocortex and maintain proliferative capability (Dawson et al., 2003; Gensert and Goldman, 2001; Nunes et al., 2003). Therefore, three successive stages of oligodendrocytes are often found: OPCs, immature/non-myelin-forming oligodendrocytes (IMO), and mature/myelin-forming oligodendrocytes (MO) (Silbereis et al., 2010). While accumulating evidence suggest that glial cells in the neocortex are diverse in properties and function (Bayraktar et al., 2020; Marques et al., 2016; Zeisel et al., 2018), little is known about the developmental origin of the diversity, especially at the progenitor level.

To address the origin and diversity of glial cell production in the developing neocortex, it is necessary to perform quantitative lineage analyses of individual RGPs in vivo with regard to their proliferative activity, potency and glial cell output at single cell resolution. Mosaic Analysis with Double Markers (MADM) represents a powerful and unique approach for assessing native stem or progenitor cell division pattern and progeny output in a spatiotemporally controllable manner (Bonaguidi et al., 2011; Gao et al., 2014; Hippenmeyer et al., 2010; Mihalas and Hevner, 2018; Zong et al., 2005). To determine RGP behavior and the cellular programs underlying neocortical gliogenesis, we exploited the unprecedented resolution of MADM and performed a systematic and quantitative clonal analysis of gliogenesis in the mouse neocortex. We found that RGPs undergo the transition from neurogenesis into gliogenesis progressively, with a peak at embryonic day (E) 16. Upon transition, individual RGPs proceed to generate astrocytes, oligodendrocytes, or both in the proportion of ∼60%:15%:25%, respectively. Notably, the generation of astrocytes and oligodendrocytes appears to be independent at the level of individual RGPs. Clonally related astrocytes or oligodendrocytes form discrete, unmixed local clusters, consistent with the existence of fate-restricted intermediate astrocyte or oligodendrocyte precursor cells originating from individual gliogenic RGPs. While the generation of fate-restricted intermediate precursor cells appears to be stochastic, individual precursor cells display clear patterns in number and subtype of glia that they generate. Interestingly, clonal removal of Neurofibromatosis type 1 (NF1), a tumor suppressor protein (Ratner and Miller, 2015), leads to excessive production of glial cells, especially oligodendrocyte precursor cells, but not neurons, indicating the unique susceptibility of the oligodendrocyte lineage in brain tumor formation. Together, these results define in vivo RGP behavior and the cellular program of gliogenesis in the mammalian neocortex quantitatively and suggest the distinct lineage origin of primary brain tumorigenesis.

## RESULTS

### MADM analysis of neocortical gliogenesis

To label selectively RGPs in the developing mouse neocortex in a temporal-specific manner, we introduced the *Emx1-CreERT2* transgene (Kessaris et al., 2006) into the MADM system (Gao et al., 2014; Hippenmeyer et al., 2010) **(Figure 1A)**. In MADM, Cre recombinase-mediated interchromosomal recombination in the G_2_ phase of dividing progenitors followed by X-segregation (G_2_-X, segregation of recombinant sister chromatids into separate daughter cells) reconstitutes one of two fluorescent markers, enhanced green fluorescence protein (EGFP, green) or tandem dimer Tomato (tdTomato, red), in each of the two daughter cells **(Figure S1A)** (Zong et al., 2005). As such, G_2_-X MADM events result in permanent and distinct labeling of the two descendent lineages, thereby allowing a direct assessment of the division pattern (symmetric versus asymmetric) and potential (the number of progeny) of the original dividing progenitors. In addition, upon G_2_-Z (congregation of recombinant sister chromatids into the same daughter cell), G_1_, or G_0_ recombination events, green and red fluorescent proteins are restored simultaneously in the same cell, resulting in double-labeled (yellow) cells **(Figure S1A)**.

**Figure 1.**
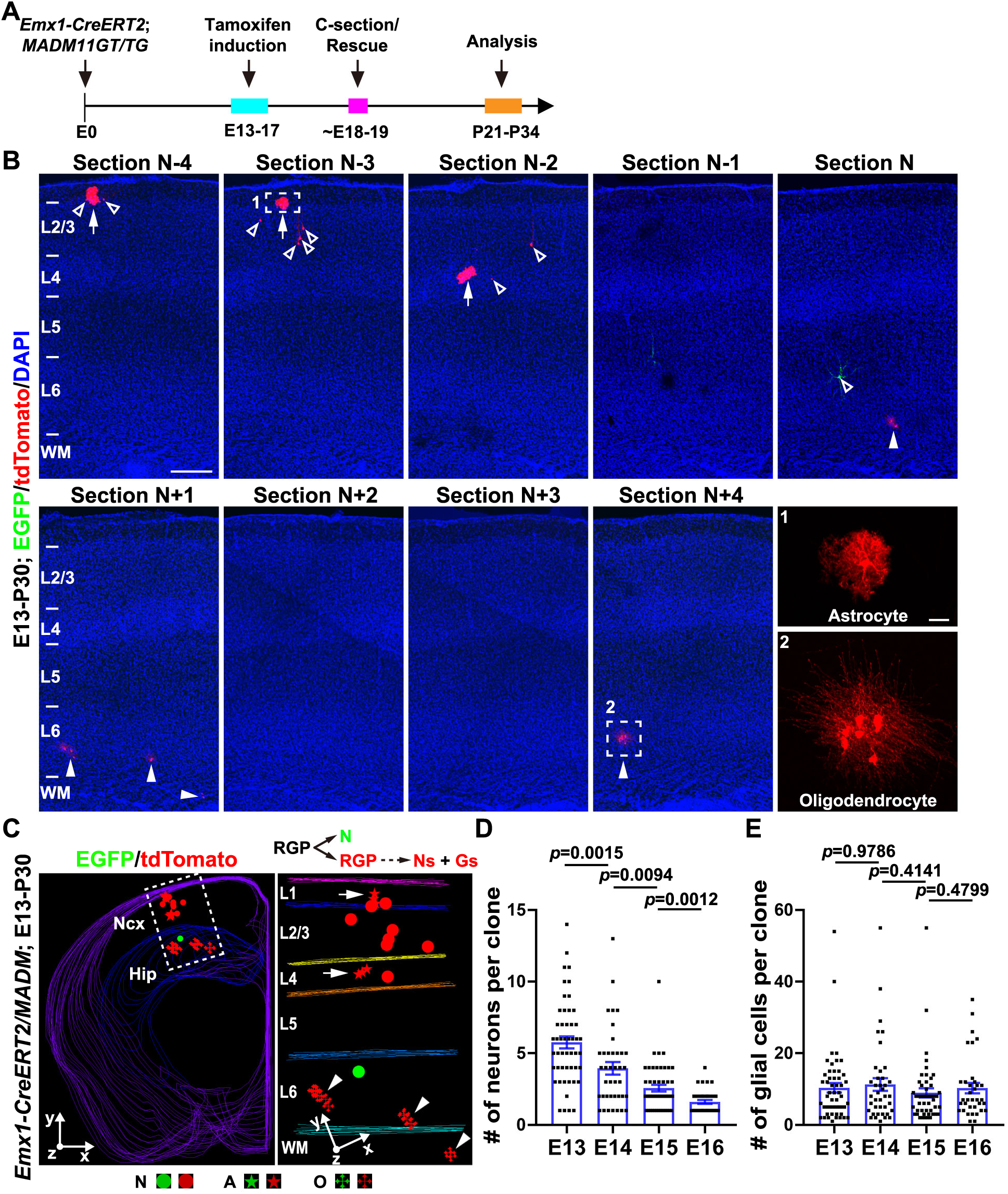
**MADM labeling of gliogenic clones originated from RGPs in the developing neocortex.** (A) Experimental paradigm of MADM-based clonal analysis. (B) Confocal images of a representative green/red fluorescence G_2_-X clone containing neurons and glial cells (N + G) labeled at E13 and examined at P30 (E13-P30). Consecutive brain sections were stained with antibodies against EGFP (green) and tdTomato (red), and with DAPI (blue). Layers are shown to the left. Arrows indicate astrocytes, arrowheads indicate oligodendrocytes, and open arrowheads indicate excitatory neurons. High-magnification images of an astrocyte (broken line, area 1) and a few oligodendrocytes (broken line, area 2) are shown in insets. Scale bars: 200 μm and 20 μm. (C) 3D reconstruction image of the E13-P30 green/red fluorescence G_2_-X N + G clone shown in B. Zoom-in image of the clone (broken line) is shown to the right. The division mode and progeny output of the originally labeled RGP are shown at the top. Colored lines indicate the brain contours and layer boundaries. Round dots indicate the cell bodies of labeled neurons. Stars indicate the cell bodies of labeled astrocytes and crosses indicate the cell bodies of labeled oligodendrocytes. The x/y/z axes indicate the spatial orientation of the clone with the y axis parallel to the brain midline and pointing dorsally. Arrows indicate the discrete astrocyte subclusters and arrowheads indicate the discrete oligodendrocyte subclusters. Note that in the entire hemisphere, only one G_2_-X N + G clone is labeled, and that clonally labeled astrocytes and oligodendrocyte form discrete subclusters nearby clonally related neurons. Ncx, neocortex; Hip, Hippocampus; L, layer; WM, white matter; RGP, radial glial progenitor; N, neuron; G, glia; A, astrocyte; O, oligodendrocyte. Similar style and labels are used in subsequent figures. (D) Quantification of the number of neurons in individual G_2_-X N + G clones labeled at different embryonic stages (E13, n=50; E14, n=39; E15, n= 48; E16, n=35). Note the progressive decrease in the number of neurons in individual clones as the labeling time proceeds. Bar plots and lines represent mean ± SEM and dots represent individual clones. Statistical analysis was performed using two-sided Mann-Whitney-Wilcoxon test. (E) Quantification of the number of glial cells in individual G_2_-X N + G clones labeled at different embryonic stages (E13, n = 50; E14, n = 39; E15, n = 48; E16, n = 35). Note the number of glial cells in individual clones remains largely constant across different labeling time points. Bar plots and lines represent mean ± SEM and dots indicate individual clones. Statistical analysis was performed using two-sided Mann-Whitney-Wilcoxon test.

Previous studies showed that RGPs switch from an amplification phase of symmetric division to a neurogenic phase of asymmetric division around E12 (Gao et al., 2014). To assess the glial cell output of a single RGP on entry into its gliogenic phase, we therefore induced Cre activity through a single dose of tamoxifen (TM) administered to timed pregnant females at one of the following embryonic stages: E13, E14, E15, E16 and E17. To ensure strict clonal density labeling, we titrated the TM dose to achieve very sparse labeling (see below). Brains were analyzed at postnatal day (P) 21-34, when gliogenesis in the neocortex is largely complete (Reemst et al., 2016). To recover all labeled cells in the neocortex, we performed serial sectioning and three-dimensional (3D) reconstruction of individual brains **(Figure 1B and 1C)**.

As expected, we observed cells labeled in green (EGFP) or red (tdTomato) fluorescence (G_2_-X), or both colors (G_2_-Z or G_1/0_ recombination, yellow), which formed spatially distinct clusters **(Figure 1B)**. Labeled cells included neurons and glial cells that could be readily distinguished based on their characteristic morphological features and specific marker expression **(Figure S1B-D)**. Neurons typically possessed a pyramid-shaped cell body and long neurites, and expressed the neuronal marker NEUN **(Figure S1B)**. In contrast, glial cells usually contained a round shape cell body and grew numerous short processes **(Figure S1C and S1D)**. Moreover, astrocytes often possessed highly branched bushy processes (PA) or a ‘star-like’ appearance (FA), and expressed S100β **(Figure S1C)**, whereas oligodendrocytes typically developed parallel processes and expressed OLIG2 **(Figure S1D)**. We selectively focused on G_2_-X green or red fluorescent cell clusters, as they represented definitively individual clones derived from the labeling of a single dividing RGP. Moreover, we singled out G_2_-X clones containing both neurons and glial cells, as these clones must have undergone the transition from neurogenesis to gliogenesis and thereby constitute the full gliogenic output of individual RGPs as they enter the gliogenic phase. To distinguish explicitly between astrocytes and oligodendrocytes, we stained the brain sections with an antibody against OLIG2 **(Figure S1D)**.

Figure 1B and 1C show a typical clone from a P30 brain treated with TM at E13 (E13-P30) **(Movie S1)**. In this case, in the entire hemisphere, only one green or red G_2_-X clonal cluster was observed with both neurons and glial cells, including astrocytes and oligodendrocytes. With one green fluorescent neuron (“minority”) and the other neurons and glial cells (“majority”) labeled in red, this clone likely derives from a labeled RGP that underwent an asymmetric neurogenic division to produce a deep layer neuron (green) and a self-renewing RGP (red) that subsequently went through multiple rounds of division to produce a number of neurons as well as glia **(Figure 1C, top inset)**. Notably, labeled glial cells were located in close proximity to their clonally related neurons, forming discrete subclusters **(Figure 1C, arrows and arrowheads)**, an arrangement confirmed by nearest-neighbor distance (NND) analysis (Brown et al., 2011; Diggle, 2003; Gao et al., 2014) of cell body positions **(Figure S2A)**. Similar observations have been reported previously (Clavreul et al., 2019; Gao et al., 2014; Garcia-Marques and Lopez-Mascaraque, 2013; Magavi et al., 2012). Consistent with previous studies (Clavreul et al., 2019; Foo et al., 2011), we did not find evidence for extensive apoptosis during neocortical gliogenesis, based on the expression of cleaved Caspase 3 **(Figure S2B)**.

We then analyzed systematically the ensemble of G_2_-X neuron- and glial cell-containing (i.e., N + G) clones labeled at E13, E14, E15 and E16. Interestingly, while the average number of neurons in individual clones decreased gradually with progressively later labeling time **(Figure 1D)**, consistent with the progressive restriction in neocortical neurogenesis over time, the average number of glial cells in individual clones remained largely constant across different labeling time points **(Figure 1E)**. On average, following entry into their gliogenic phase, individual glial cell-producing RGPs produced ∼10 glial cells, including astrocytes and/or oligodendrocytes. These results suggest that individual gliogenic RGPs generate a similar number of glial cells regardless of the labeling time at E13-E16. Moreover, the number of glial cells in individual clones located in different neocortical areas was largely comparable **(Figure S2C-E)**, suggesting that the gliogenic capability is a conserved feature of RGPs across the developing neocortex.

### A defined pattern of RGP gliogenesis

We next analyzed glial cell composition of individual G_2_-X N + G clones labeled at different embryonic stages. We consistently observed three types of clones: neuron plus astrocyte-containing (i.e., N + A; **Figure 2A left and S3**), neuron plus oligodendrocyte-containing (i.e., N + O; **Figure 2A middle and S4**), and neuron plus astrocyte and oligodendrocyte-containing (i.e., N + A + O; **Figure 1 and 2A right)**. Moreover, the relative fraction of N + A, N + O, and N + A + O clones labeled at different embryonic stages was largely consistent across different induction times, with ∼60% of gliogenic RGPs producing only astrocytes, ∼15% producing only oligodendrocytes, and ∼25% producing both astrocytes and oligodendrocytes **(Figure 2B and 2C)**. These results suggest that, upon transition into the gliogenic phase, RGPs exhibit a defined propensity in producing astrocytes and/or oligodendrocytes. While the majority (∼75%) of gliogenic RGPs produce only astrocytes or oligodendrocytes, a significant portion (∼25%) gives rise to both astrocytes and oligodendrocytes, indicating the existence of bipotent glial progenitors in the embryonic neocortex, as previously suggested in the embryonic spinal cord and neonatal optic nerve (Herrera et al., 2001; Raff et al., 1983).

**Figure 2.**
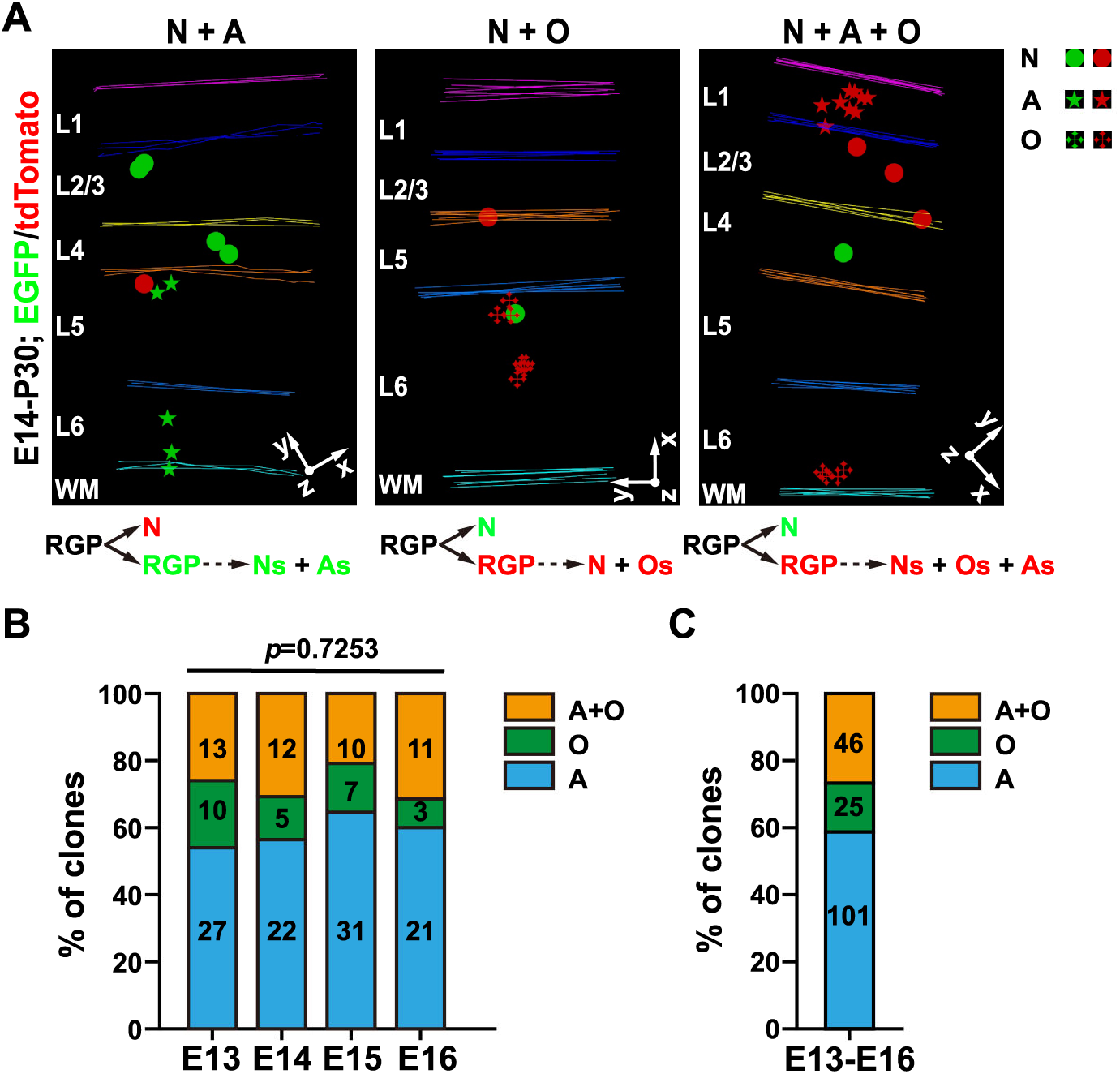
**Three different types of gliogenic clones.** (A) 3D reconstruction images of three representative types of gliogenic G_2_-X N + G clones labeled at E14 and examined at P30 (E14-P30), including astrocyte only (N + A, left), oligodendrocyte only (N + O, middle), and astrocyte and oligodendrocyte (N + A + O, right). The division modes and progeny output of the originally labeled RGPs are shown at the bottom. Layers are shown to the left. Colored lines indicate the brain contours and layer boundaries. The x/y/z axes indicate the spatial orientation of the clone with the y axis parallel to the brain midline and pointing dorsally. (B) Quantification of the relative fraction of N + A clone (A), N + O clone (O), and N + A + O clone (A+O) labeled at different embryonic stages. Note the similar rate of three different types of clones across different labeling time points. Statistical analysis was performed using Chi-square test. (C) Quantification of the overall fraction of three different types of gliogenesis clones (A, ∼60%; O, ∼15%; A + O, ∼25%) labeled at E13-16.

### Three distinct modes of gliogenesis transition

MADM labeling allows the division mode of RGPs to be inferred explicitly from the fluorescence color and respective cell fate of the progeny. Thus, it is possible to capture directly RGPs that are actively transitioning from neurogenesis to gliogenesis (i.e., RGPs undergoing the last neurogenic division to proceed to gliogenesis) as the labeling occurs **(Figure 3A, top)**. Such events, resulting in neuron(s) in one color and glial cell(s) in the other color, were indeed observed among the G_2_-X clones **(Figure 3A, middle)**. As expected, we also observed prior-transitioning N + G clones with neurons in both colors (i.e., where labeling occurred prior to the onset of gliogenesis; **Figure 3A, left**), and glial cell-only clones (i.e., where labeling occurred after the onset of gliogenesis; **Figure 3A, right**). Notably, very few transitioning clones were observed at E13 **(Figure 3B)**. The fraction of transitioning clones progressively increased and peaked at E16. The vast majority of clones labeled at E17 contained only glial cells **(Figure S5A)**, indicating that RGPs have largely transitioned to gliogenesis by E17.

**Figure 3.**
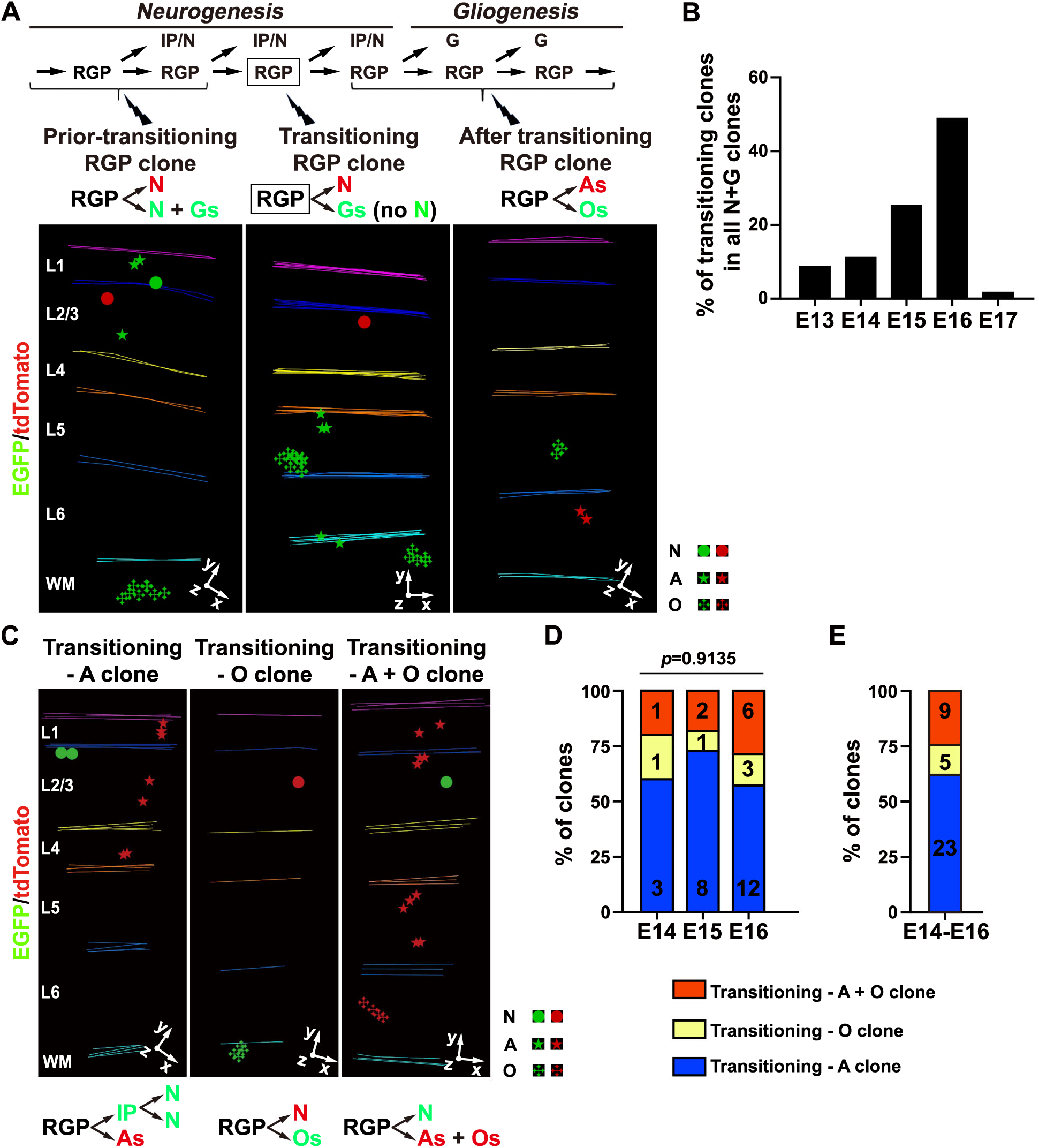
**RGPs transition to gliogenesis in three different modes.** (A) 3D reconstruction images of representative G_2_-X labeled prior-transitioning RGP clone (left), transitioning RGP clone (middle), and after transitioning RGP clone (right). The lineage progression, division mode and progeny output of RGPs are shown at the top. Thunder shapes indicate MADM labeling of RGPs and rectangles indicate the transitioning RGP. IP, intermediate progenitor; N, neuron; G, glia. Note that transitioning clone contains neurons and glial cells in different colors, whereas prior-transitioning clone contains neurons and glial cells in the same color. After transitioning clones contain glia cells only. (B) Quantification of the fraction of transitioning clones at different embryonic stage. Note that RGPs transition to gliogenesis largely at E16. (C) 3D reconstruction images of three representative gliogenesis transition clones, including transitioning-A clone (left), transitioning-O clone (middle), and transitioning-A + O clone (right). The division mode and progeny output of RGPs are shown at the bottom. (D) Quantification of the relative fraction of three different gliogenesis transition modes at different embryonic stages. Note the consistent frequencies of three different gliogenesis transitioning modes across different embryonic stages. Statistical analysis was performed using Chi-square test. (E) Quantification of the overall fraction of three gliogenic transitioning modes (A, ∼60%; O, ∼15%; A + O, ∼25%) observed at E14-E16.

Consistent with the three types of gliogenic clone (N + A, N + O, and N + A + O), we observed three corresponding types of transitioning RGP clones **(Figure 3C)** suggesting that, upon transitioning to gliogenesis, individual RGPs enter one of the three modes of gliogenesis to produce astrocytes, oligodendrocytes, or both. Remarkably, the relative proportion of these three gliogenic transition modes was largely similar across different embryonic stages **(Figure 3D)**. That is, the rate of RGP transitioning to produce astrocytes, oligodendrocytes, or both was ∼60%:15%:25%, respectively **(Figure 3D and 3E)**, the same as the relative fraction of three types of gliogenesis clone **(Figure 2B and 2C)**. These results suggest that gliogenic transition and subsequent gliogenesis mode is a conserved and predictable feature of RGPs in lineage progression, regardless of the exact timing of transition.

### Independent generation of astrocyte and oligodendrocyte

Having found that individual RGPs may generate astrocytes, oligodendrocytes, or both glial cell types, we tested whether the generation of astrocytes or oligodendrocytes influences each other at the single RGP level. We compared the number of astrocytes between N + A and N + A + O clones **(Figure S5B)**. Interestingly, the number of astrocytes in N+A clones was similar to that in N + A + O clones **(Figure S5C)**, suggesting that astrocyte generation by individual RGPs is not affected by whether the same RGPs produce oligodendrocytes. Similarly, the number of oligodendrocytes in N + O clones was comparable to that in N + O + A clones **(Figure S5D and S5E)**, indicating that oligodendrocyte generation by individual RGPs is not affected by whether the same RGPs produce astrocytes. Together, these results suggest that astrocyte and oligodendrocyte generation by individual RGPs is largely independent of each other.

Given that ∼25% of gliogenic RGPs produce both astrocytes and oligodendrocytes, the independence of astrocyte and oligodendrocyte generation predicts that the total glial cell output in N + A + O clones would be close to the sum of the glial output of N + A and N + O clones. Indeed, this was the case **(Figure S5F)**. These results further suggest that individual RGPs generate astrocytes and oligodendrocytes as two independent processes.

### Local glial generation by fate-restricted precursor cells

To further understand the cellular processes underlying gliogenesis by RGPs, we systematically analyzed glial cell number and spatial organization of the clone. Notably, within individual clones, astrocytes and oligodendrocytes were often localized in discrete subclusters **(Figure 1B, 1C, 2A, 3A, 3C, S5B and S5D)**, raising the possibility that individual subclusters may arise from local precursor cells originating from RGPs. Mitotic glial precursor cells have been shown to exist in the neonatal cortex (Ge et al., 2012; Levison and Goldman, 1993; Parnavelas, 1999) and are positive for NESTIN (Siddiqi et al., 2014). To test this directly, we introduced the *Nestin-CreERT2* transgene (Imayoshi et al., 2006) to the MADM system and induced Cre activity at P3 or P5. As expected, we observed green or red, as well as yellow, fluorescent glial cells, including either astrocytes or oligodendrocytes, which formed spatially-isolated clusters **(Figure 4A, S6A, and S6B)**.

**Figure 4.**
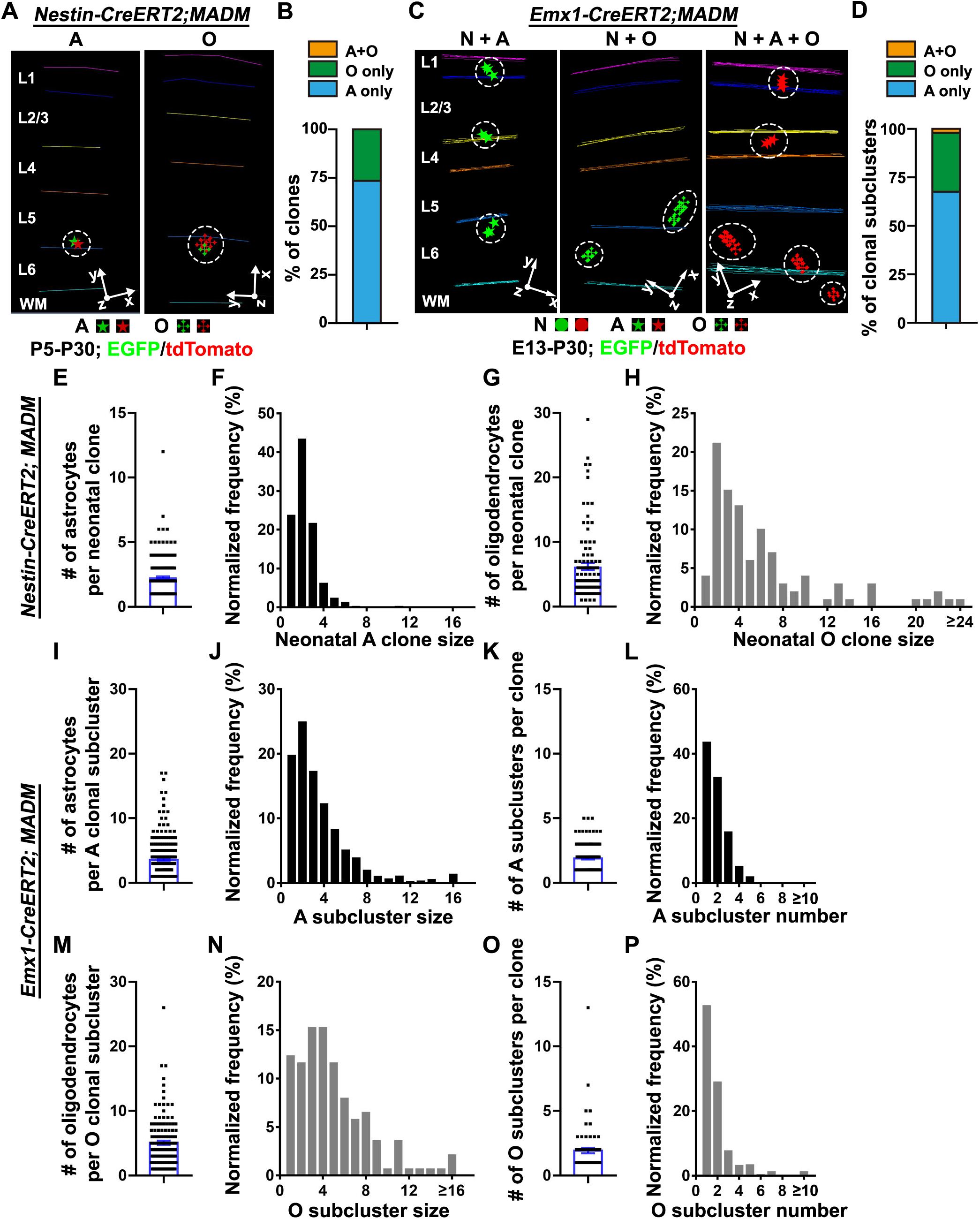
**RGPs generate glial cells via fate-restricted precursor cells that produce local clusters.** (A) 3D reconstruction images of representative G_2_-X clones labeled at P5 and examined at P30 in the *Nestin-CreERT2;MADM* neocortex. Note that individual clonal clusters (broken lines) contained only astrocyte (A, left) or oligodendrocyte (O, right). (B) Quantification of the glial cell composition (astrocyte and oligodendrocyte) of individual clones labeled at the neonatal stage in the *Nestin-CreERT2;MADM* neocortex (n = 380). Note that individual neonatal clones contain either astrocyte or oligodendrocyte, but not both, indicating that they are originated from fate restricted astrocyte or oligodendrocyte precursor cells. (C) 3D reconstruction images of representative E13-P30 G_2_-X N + A, N +O, and N + A + O clones in the *Emx1-CreERT2;MADM* neocortex consisting of discrete glial cell subclusters (broken lines) identified using mean shift analysis. Neurons are not shown in the images. (D) Quantification of the glial cell composition (astrocyte and oligodendrocyte) of individual subclusters in clones labeled at the embryonic stage in the *Emx1-CreERT2;MADM* neocortex (n = 400). Note that nearly all individual subclusters contain either astrocyte or oligodendrocyte, but not both, indicating that they are derived from fate-restricted astrocyte or oligodendrocyte precursor cells. (E) Quantification of the number of astrocyte in individual neonatal astrocyte clones (n = 273). Bar plots and lines represent mean ± SEM and dots represent individual clones. (F) Histogram of the number of astrocyte in individual neonatal astrocyte clones (n = 273). Note that the distribution displays a peak, indicating a relatively defined size in astrocyte precursor cell output. (G) Quantification of the number of oligodendrocyte in individual neonatal oligodendrocyte clones (n = 107). Bar plots and lines represent mean ± SEM and dots represent individual clones. (H) Histogram of the number of oligodendrocyte in individual neonatal oligodendrocyte clones (n = 107). Note that the distribution displays a peak, indicating a relatively defined size in oligodendrocyte precursor cell output. (I) Quantification of the number of astrocyte in individual astrocyte subclusters (n = 273) identified in the embryonic clones. Bar plots and lines represent mean ± SEM and dots represent individual subclusters. (J) Histogram of the number of astrocyte in individual astrocyte subclusters (n = 273). Note that the distribution displays a peak, indicating a relatively defined size of the astrocyte subcluster. (K) Quantification of the number of astrocyte subcluster in individual embryonic clones (n = 147). Bar plots and lines represent mean ± SEM and dots represent individual clones. (L) Histogram of the number of astrocyte subcluster in individual embryonic clones (n = 147). (M) Quantification of the number of oligodendrocyte in individual oligodendrocyte subclusters (n = 136) identified in the embryonic clones. Bar plots and lines represent mean ± SEM and dots represent individual subclusters. (N) Histogram of the number of oligodendrocyte in individual oligodendrocyte subclusters (n = 136). Note that the distribution displays a peak, indicating a relatively defined size of the oligodendrocyte subcluster. (O) Quantification of the number of oligodendrocyte subcluster in individual embryonic clones (n = 71). Bar plots and lines represent mean ± SEM and dots represent individual clones. (P) Histogram of the number of oligodendrocyte subcluster in individual embryonic clones (n = 71).

We then focused on G_2_-X green or red fluorescent neonatal glial cell clones **(Figure 4A, S6A, and S6B)**. Notably, individual clones existed predominantly as a single cluster and contained only astrocytes or oligodendrocytes, but not both **(Figure 4A and 4B)**, consistent with them being derived from a single dividing astrocyte or oligodendrocyte precursor cell. These results also suggest that the precursor cells are not bipotent, but fate-restricted to generate only astrocytes or oligodendrocytes. Moreover, the average number of astrocytes or oligodendrocytes (both green and red fluorescent) in individual clones labeled at P3 and P5 was largely the same **(Figure S6C and S6D)**, indicating that individual astrocyte or oligodendrocyte precursor cells have a relatively consistent output regardless of the labeling time. In particular, individual astrocyte precursor cells produced ∼2-3 astrocytes on average **(Figure 4E)**, while individual oligodendrocyte precursor cells produced ∼6 oligodendrocytes on average **(Figure 4G)**. In spite of the clear variability, the distribution of astrocyte or oligodendrocyte numbers in individual neonatal clones exhibited a clear peak **(Figure 4F and 4H**), indicating that individual astrocyte or oligodendrocyte precursor cells have a relatively restricted output. The average number of oligodendrocytes in individual neonatal clones was significantly larger than that of astrocytes **(Figure S6E)**, suggesting that oligodendrocyte precursor cells possess more division potential with a larger output than astrocyte precursor cells. Together, these results suggest that fate-restricted glial precursor cells divide in the neonatal neocortex to produce a relatively defined number of astrocytes or oligodendrocytes forming local clusters.

Glial cells in the embryonic RGP clones labeled using *Emx1-CreERT2* were localized predominantly in subclusters **(Figure 4C)**, suggesting that during gliogenesis, RGPs give rise to spatially segregated precursor cells. To assess this quantitatively, we performed the mean-shift clustering analysis with spatial parameters informed by the distribution of neonatal precursor cell clones to identify local subclusters within individual embryonic RGP clones **(Figure S7)**. Interestingly, we found that individual subclusters identified in embryonic RGP clones consisted of either astrocytes or oligodendrocytes, but not both **(Figure 4D)**, consistent with the notion that they originate from individual fate-restricted precursor cells. The existence of multiple astrocyte and/or oligodendrocyte subclusters in some clones indicates that individual gliogenic RGPs are capable of generating multiple astrocyte and/or oligodendrocyte precursor cells, likely via asymmetric cell division.

We then examined quantitatively the number of astrocyte or oligodendrocyte subclusters (i.e., subcluster number) in individual embryonic RGP clones as well as the number of astrocytes or oligodendrocytes (i.e., subcluster size) in individual subclusters **(Figure 4I-P)**. The average number of astrocytes in individual astrocyte subclusters (i.e., A subcluster size) was ∼3 **(Figure 4I)**, whereas the average number of oligodendrocytes in individual oligodendrocyte subclusters (i.e., O subcluster size) was ∼5-6 **(Figure 4M)**. Moreover, the histogram of A and O subcluster sizes exhibited a peak **(Figure 4J and 4N)**. These results are highly consistent with the behavior and output of astrocyte or oligodendrocyte precursor cells labeled at the neonatal stage **(Figure 4E-H)**, supporting the conclusion that astrocyte or oligodendrocyte subclusters in embryonic RGP clones are the progeny of individual astrocyte or oligodendrocyte precursor cells generated by gliogenic RGPs.

The average number of astrocyte or oligodendrocyte subclusters (i.e., A or O subcluster number) in individual embryonic RGP clones was ∼2 **(Figure 4K and 4O)**; yet, the histogram of A and O subcluster numbers in individual clones could be well approximated by a geometric distribution **(Figure 4L, 4P, S8A, S8B)**, suggesting that the number of astrocyte or oligodendrocyte precursor cells generated by individual gliogenic RGPs is stochastic. Together, these results suggest that individual gliogenic RGPs generate a stochastic number of astrocyte or oligodendrocyte precursor cells, which in turn produce ∼2-3 astrocytes or ∼5-6 oligodendrocytes, respectively, forming a local subcluster.

### Subtype output of individual gliogenic RGPs

Previous studies described three major subtypes of astrocytes in the neocortex: L1A, PA and FA, which can be readily distinguished based on their morphological features, location and marker expression (Garcia-Marques and Lopez-Mascaraque, 2013; Middeldorp and Hol, 2011; Zeisel et al., 2018). Indeed, we observed three subtypes of astrocyte with distinct morphological features and GFAP expression level **(Figure 5A)**. We systematically analyzed the subtype composition of individual embryonic RGP clones. More than 60% of RGP clones with astrocytes contained only one subtype of astrocyte **(Figure 5B)**, predominantly PA (∼70%) or FA (∼27%) **(Figure 5D)**. About 30% of RGP clones with astrocytes contained two subtypes of astrocyte, either PA and FA (PA + FA) or L1A and PA (L1A + PA), in a similar occurrence **(Figure 5C)**. Only a very small fraction (3%) contained all three subtypes (L1A + PA + FA). We did not observe any RGP clones containing FA (predominantly located in the white matter) and L1A (located in layer 1). These results suggest that astrocyte generation by individual RGPs exhibits layer and subtype-specific regulation.

**Figure 5.**
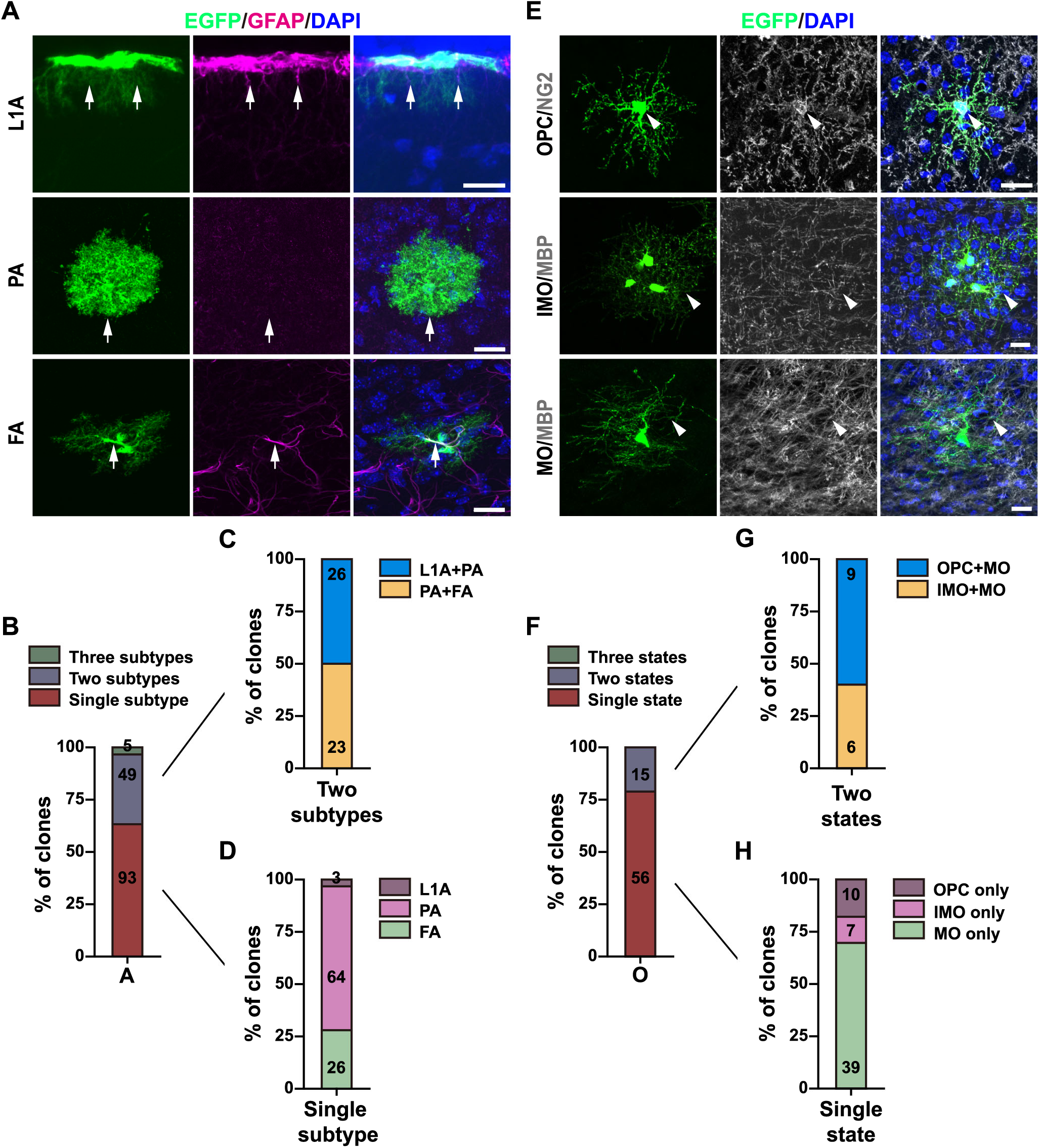
**Subtype organization of glial cells in RGP clones.** (A) Confocal images of MADM labeled astrocytes stained with antibodies against EGFP (green) and GFAP (magenta), and with DAPI (blue). L1A, L1 astrocyte; PA, protoplasmic astrocyte; FA, fibrous astrocyte. Note the three distinct subtypes of astrocyte reflected in morphology and/or marker expression. Scale bars, 20 μm. (B) Quantification of the fraction of embryonic RGP clones containing single, two, or three subtypes of astrocyte. (C) Quantification of the subtype organization of embryonic RGP clones with two different subtypes of astrocyte. (D) Quantification of the subtype organization of embryonic RGP clones with single subtype of astrocyte. (E) Confocal images of MADM labeled oligodendrocytes stained with antibodies against EGFP (green) and NG2 (white, top), or MBP (white, middle and bottom), and with DAPI (blue). Note the three distinct differentiation states of oligodendrocyte reflected in morphology and/or marker expression. OPC, oligodendrocyte precursor cell; IMO, immature oligodendrocyte; MO, mature oligodendrocyte. Scale bars, 20 μm. (F) Quantification of the fraction of embryonic RGP clones containing single, two, or three differentiation states of oligodendrocyte. (G) Quantification of the differentiation state organization of embryonic RGP clones with two different differentiation states of oligodendrocyte. (H) Quantification of the differentiation state organization of embryonic RGP clones with single different differentiation state of oligodendrocyte.

To further test this, we examined the subtype composition of individual astrocyte subclusters in embryonic RGP clones. Interestingly, individual astrocyte subclusters contained predominantly the same subtype of astrocyte, including PA (∼65%), FA (∼20%) and L1A (∼5%), with the remainder mixed **(Figure S8C)**. These results suggest that individual astrocyte precursor cells generate mostly the same subtype of astrocyte.

Previous studies suggest that neocortical oligodendrocytes can be categorized into three differentiation or maturation states, OPC, IMO and MO, as reflected in their morphological features and molecular expression (Miller and Ono, 1998; Zhang, 2001). Consistent with previous studies, we observed NG2-expressing OPCs with thick and bushy branches, IMO with thin and bushy branches, and Myelin Basic Protein (MBP)-expressing MO with parallel branches **(Figure 5E)**. Interestingly, the majority (>75%) of embryonic RGP clones with oligodendrocytes contained oligodendrocytes in only one differentiation state **(Figure 5F)**, including MO (∼70%), OPC (∼20%), or IMO (∼10%) **(Figure 5H)**. About 20% of these embryonic RGP clones contained oligodendrocytes in two differentiation states, OPC and MO (OPC + MO, ∼60%) or IMO and MO (IMO + MO, ∼40%) **(Figure 5G)**. No embryonic RGP clones contained oligodendrocytes in all three differentiation states (OPC + IMO + MO) or OPC and IMO (OPC + IMO). These results suggest that oligodendrocyte generation by RGPs in the developing neocortex is regulated. Notably, the vast majority of oligodendrocyte subclusters in embryonic RGP clones contained oligodendrocytes in only one differentiation state, including MO (∼68%), OPC (∼17%) or IMO (∼10%) **(Figure S8D)**. These results suggest that individual oligodendrocyte precursor cells arising from gliogenic RGPs generate predominantly oligodendrocytes with the same differentiation state.

### NF1 selectively suppresses gliogenesis

To gain insight into the molecular regulation of neocortical gliogenesis, we took advantage of the MADM design to perform single cell level loss-of-function analysis of NF1, a tumor suppressor protein that has been linked to brain tumor development (Anastasaki et al., 2017; Campian and Gutmann, 2017; Liu et al., 2011). We introduced the conditional deletion allele of *Nf1* into the *Emx1-CreERT2;MADM* system by genetically linking it to the *GT* cassette through meiotic recombination; in parallel, the *TG* cassette was linked to the wild-type allele **(Figure 6A)**. As a result, upon Cre-mediated inter-chromosomal recombination, the green fluorescent cells inherited the wild-type *Nf1* allele, whereas the red fluorescent cells inherited the *Nf1* mutant allele. Therefore, in these experiments, the red fluorescent cells represent the *Nf1* mutant cells, whereas the green fluorescent cells represent the internal wild type control cells in the same clone.

**Figure 6.**
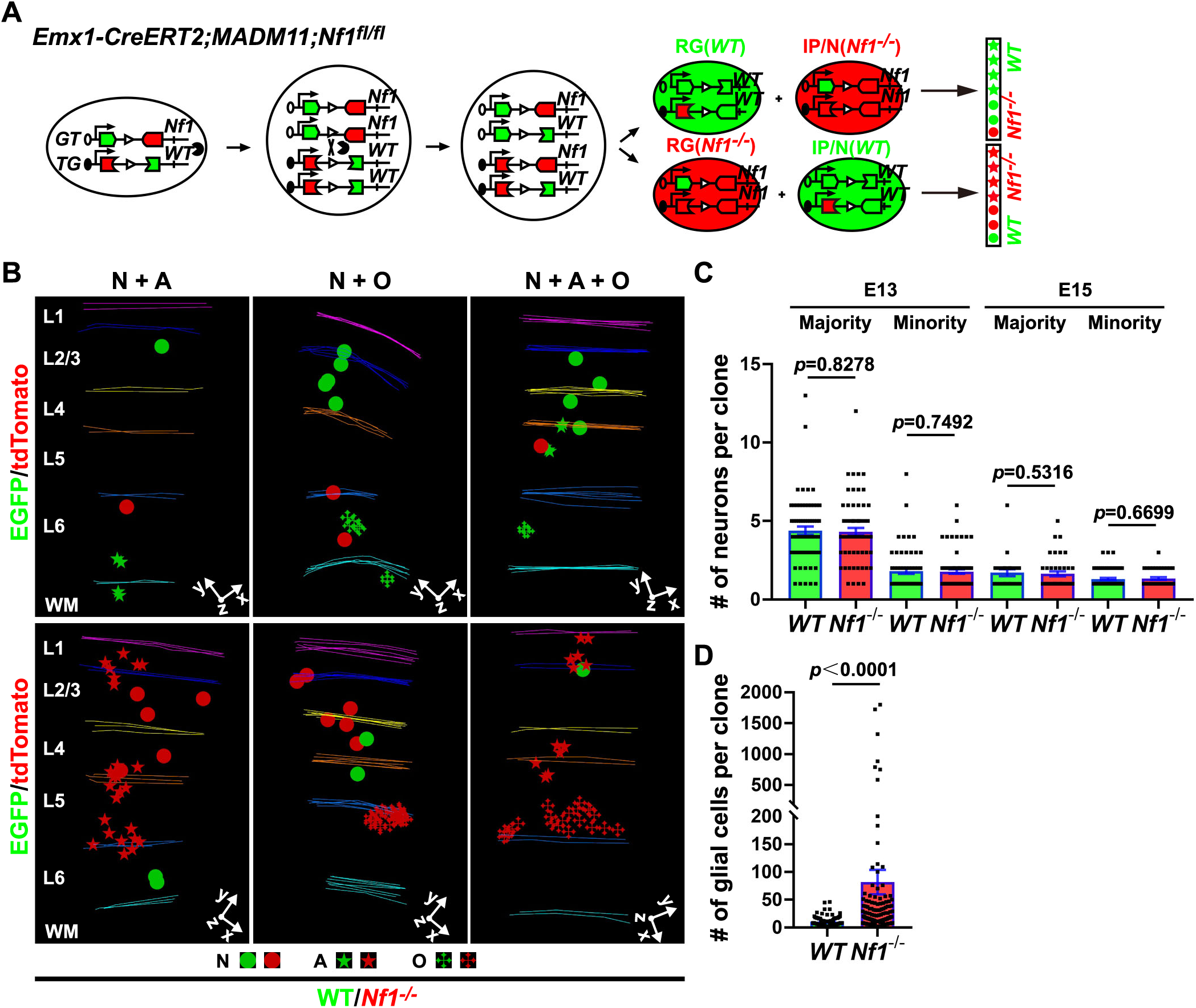
**NF1 loss selectively promotes glial cell generation.** (A) Schematics of MADM-based mosaic knockout analysis of *Nf1* in RGPs. Note that tdTomato labels *Nf1^-/-^* cells and EGFP labels wild type (WT) cells within individual clones. (B) 3D reconstruction images of representative N + A, N + O, and N + A + O clones in the *Emx1-CreERT2;MADM;Nf1^fl/fl^* neocortex labeled at E13 or E15 and analyzed at P30. Note no obvious difference between the numbers of red fluorescent *Nf1* mutant neurons as the minority or majority and those of green fluorescent WT neurons as the minority or majority in individual RGP clones; yet, the number of red fluorescent *Nf1* mutant glial cells is significantly more than that of green fluorescent WT glial cells. (C) Quantification of the number of WT or *Nf1* mutant neurons as the majority or minority in individual RGP clones (E13 majority: WT, n = 65; *Nf1^-/-^*, n = 69; E13 minority: WT, n = 61; *Nf1^-/-^*, n = 41; E15 majority: WT, n = 69; *Nf1^-/-^*, n = 41; E15 minority: WT, n = 61; *Nf1^-/-^*, n = 39). Bar plots and lines represent mean ± SEM and dots represent individual clones. Statistical analysis was performed using two-sided Mann-Whitney-Wilcoxon test. (D) Quantification of the number of WT or *Nf1* mutant glial cell number in individual RGP clones (WT, n = 107; *Nf1^-/-^*, n = 141). Bar plots and lines represent mean ± SEM and dots represent individual clones. Statistical analysis was performed using two-sided Mann-Whitney-Wilcoxon test.

We observed a similar rate of labeling in the *Emx1-CreERT2;MADM;Nf1^fl/fl^* neocortex by TM administration at E13 or E15 **(Figure 6B)**. We focused on analyzing G_2_-X green or red fluorescent clonal clusters containing both neurons and glial cells **(Figure 6B)**. Notably, the average number of red fluorescent *Nf1* mutant neurons in individual clones was the same as that of green fluorescent wild type neurons, either as the majority or minority of the clone **(Figure 6B and 6C)**, suggesting that clonal removal of NF1 in RGPs does not affect neurogenesis. In sharp contrast, the average number of red fluorescent *Nf1* mutant glial cells in individual clones was substantially larger than that of green fluorescent wild type glial cells **(Figure 6B and 6D)**, suggesting that clonal removal of NF1 strongly promotes gliogenesis by RGPs. Together, these results suggest that NF1 is not involved in the regulation of neurogenesis, but drastically suppresses gliogenesis by RGPs in the developing neocortex.

### NF1 loss causes excessive OPC generation

We further dissected the effect of NF1 removal on astrocyte and oligodendrocyte generation. Interestingly, while NF1 removal caused a significant increase in the generation of both astrocytes and oligodendrocytes, the enhancement of oligodendrocyte generation was far more drastic than that of astrocyte generation **(Figure 7A-D)**. On average, astrocyte generation exhibited ∼2-fold increase, whereas oligodendrocyte generation showed ∼15-fold increase. In particular, we observed more than 6% of red fluorescent *Nf1* mutant oligodendrocyte-containing clones that possessed hundreds or more of oligodendrocytes that formed a large cluster **(Figure 7D arrow, 7E arrows, and S8E)**. These results suggest that NF1 removal drives excessive generation of glial cells, especially oligodendrocytes.

**Figure 7.**
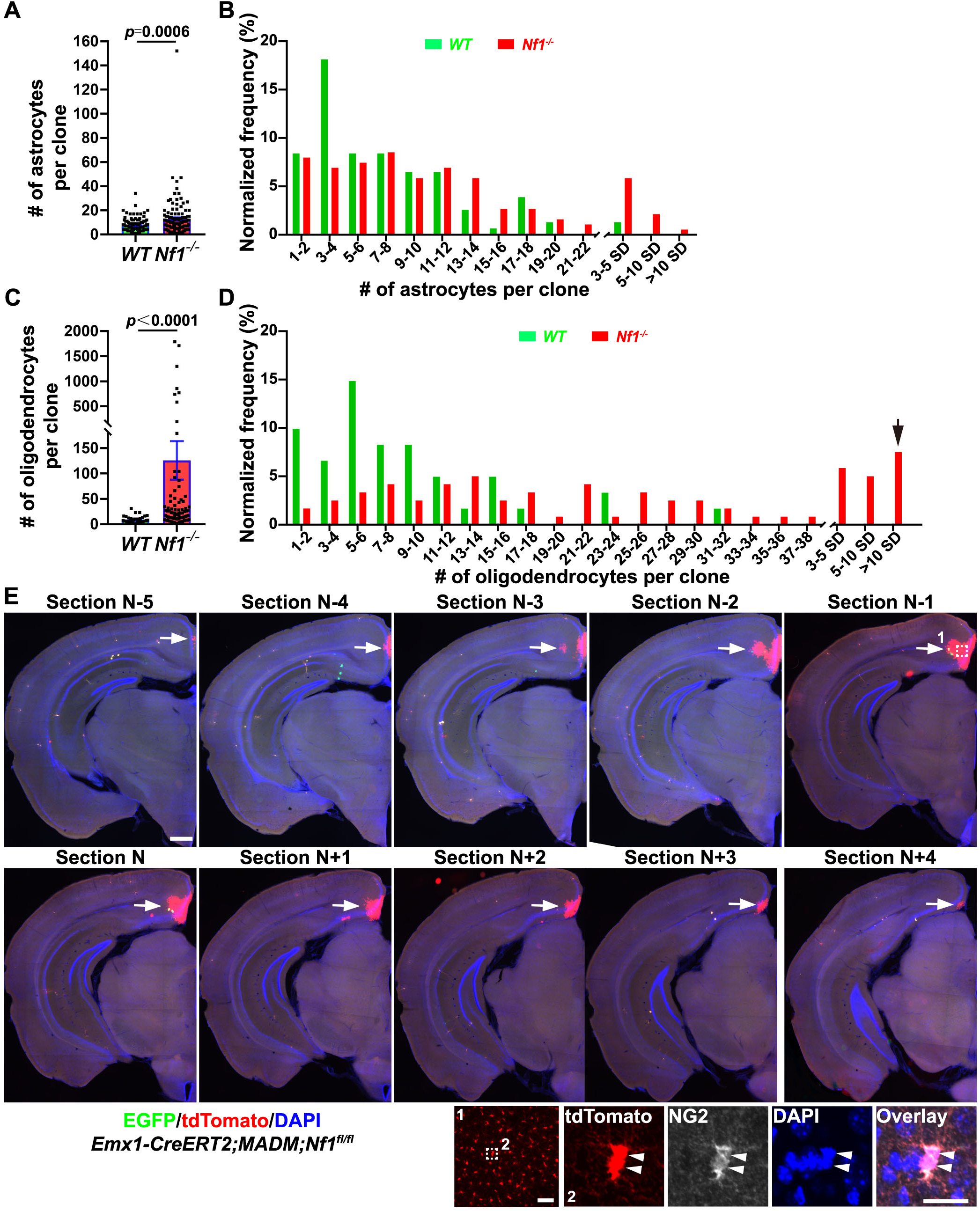
**NF1 loss causes excessive production of OPCs.** (A) Quantification of the number of WT or *Nf1^-/-^* astrocyte in individual RGP clones (WT, n = 102; *Nf1^-/-^*, n = 124). Bar plots and lines represent mean ± SEM and dots represent individual clones. Statistical analysis was performed using two-sided Mann-Whitney-Wilcoxon test. (B) Histogram of the number of WT (green) or *Nf1^-/-^* (red) astrocyte in individual RGP clones. SD, standard deviation of the mean. (C) Quantification of the number of WT or *Nf1^-/-^* oligodendrocyte in individual RGP clones (WT, n = 40; *Nf1^-/-^*, n = 79). Bar plots and lines represent mean ± SEM and dots represent individual clones. Statistical analysis was performed using two-sided Mann-Whitney-Wilcoxon test. (D) Histogram of the number of WT (green) or *Nf1^-/-^* (red) oligodendrocyte in individual RGP clones. Arrow indicates the fraction of the extraordinarily large clone with hundreds to thousands of oligodendrocyte. SD, standard deviation of the mean. (E) Confocal images of a representative G_2_-X RGP clone containing an extraordinary number of *Nf1^-/-^* OPCs in the neocortex. Zoom-in images of *Nf1^-/-^* cells (broken lines and areas 1 and 2) expressing OPC marker NG2 are shown in insets. Scale bars: 500 μm, 20 μm and 20 μm (from top to bottom).

Moreover, we found that nearly all oligodendrocytes in the extraordinarily large *Nf1* mutant clone were NG2-positive OPCs **(Figure 7E, inset)**, suggesting that NF1 removal in individual RGPs leads to the emergence of extraordinary large clones of OPCs in the neocortex, which would require extensive cell division. In comparison, while astrocyte generation was also enhanced, we did not observe any extraordinarily large clone of astrocyte lacking NF1, even though significantly more astrocyte clones were labeled. Notably, OPCs continue to divide even in adulthood (Dawson et al., 2003; Gensert and Goldman, 2001; Nunes et al., 2003). These results point to a particular susceptibility of the oligodendrocyte lineage in relation to brain tumorigenesis.

### NF1 loss leads to distinct changes in astrogenesis and oligogenesis

To further dissect the progenitor cell behavior affected by the loss of NF1, we performed the subcluster analysis of the green fluorescent wild type and red fluorescent *Nf1* mutant clones **(Figure 8)**. Interestingly, the astrocyte subcluster size was largely comparable between the wild type and *Nf1* mutant clones **(Figure 8A)**; yet, the astrocyte subcluster number in individual clones was significantly larger in *Nf1* mutant clones than that in wild type clones **(Figure 8B)**. These results suggest that NF1 loss leads to an increase in the number of astrocyte precursor cells generated by individual RGPs, but not the output of individual astrocyte precursor cells **(Figure 8C)**. In contrast, both the oligodendrocyte subcluster size and number in individual clones were significantly increased in the *Nf1* mutant clone **(Figure 8D and 8E)**. In particular, we observed a significant fraction of subclusters containing hundreds or more of oligodendrocytes. These results suggest that NF1 loss not only promotes the generation of OPCs by RGPs, but also drastically enhances the division capability and output of OPCs **(Figure 8F)**. These findings indicate that NF1 loss causes distinct changes in progenitor behavior underlying astrogenesis and oligogenesis, and that OPCs can be a crucial cellular origin of brain tumor formation.

**Figure 8.**
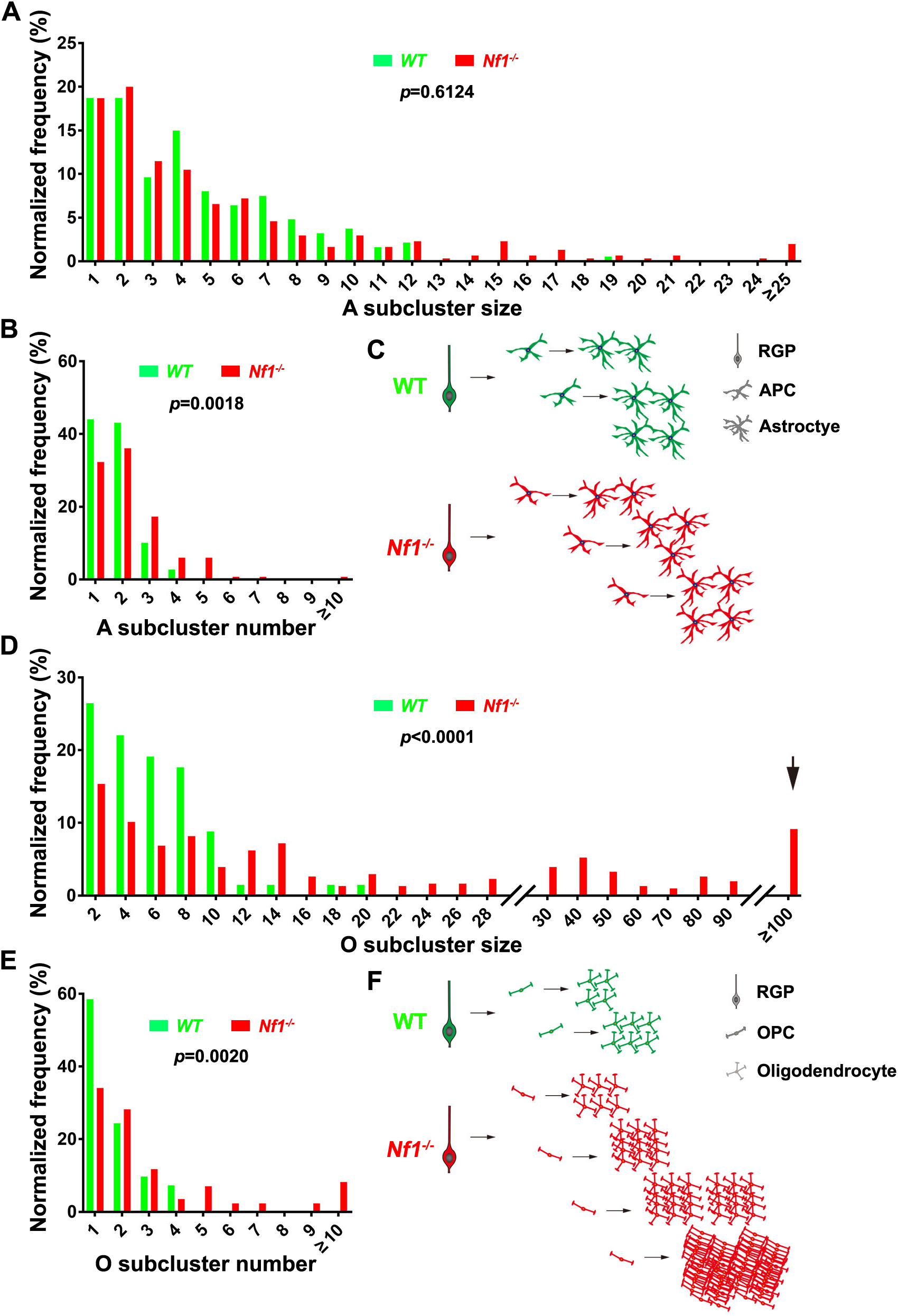
**NF1 loss leads to distinct changes in progenitor cell behaviors in astrogenesis and oligogenesis.** (A) Histogram of the numbers of astrocyte in individual WT (green, n = 187) and *Nf^1-/-^* mutant (red, n = 305) clonal subclusters. Statistical analysis was performed using two-sided Mann-Whitney-Wilcoxon test. (B) Histogram of the numbers of astrocyte subclusters in individual WT (green, n = 109) and *Nf1^-/-^* mutant (red, n = 133) clones. Statistical analysis was performed using two-sided Mann-Whitney-Wilcoxon test. (C) Schematic summary of the changes in progenitor cell behavior during astrogenesis upon NF1 loss. Note that NF1 loss leads to an increase in astrocyte precursor cell generation by RGPs, but not the output of individual astrocyte precursor cells. (D) Histogram of the numbers of oligodendrocyte in individual WT (green, n = 68) and *Nf1^-/-^* mutant (red, n = 306) clonal subclusters. Statistical analysis was performed using two-sided Mann-Whitney-Wilcoxon test. (E) Histogram of the numbers of oligodendrocyte subclusters in individual WT (green, n = 41) and *Nf1^-/-^* mutant (red, n = 85) clones. Statistical analysis was performed using two-sided Mann-Whitney-Wilcoxon test. (F) Schematic summary of the changes in progenitor cell behavior during oligogenesis upon NF1 loss. Note that NF1 loss leads to an increase in OPC generation and the output of individual OPCs; in particular, the generation of clone with hundreds or more of OPCs.

## DISCUSSION

RGPs in the developing neocortex generate neurons as well as glial cells, including astrocytes and oligodendrocytes; however, the precise behavior of RGPs and the underlying gliogenesis program remain largely unknown. In this study, we took advantage of the exquisite resolution of MADM labeling to perform a systematic and quantitative examination of neocortical gliogenesis by individual RGPs. Our in-depth analysis reveals the precise gliogenesis program at single cell resolution in the developing mouse neocortex **(Figure S8F)**. Our results show that a subset of RGPs transition from neurogenesis to gliogenesis progressively, with a peak at E16, to generate astrocytes, oligodendrocytes, or both in defined proportions of 60%:15%:25% via fate-restricted intermediate precursor cells. While the number of precursor cells generated by individual gliogenic RGPs appears to be stochastic, the output of individual precursor cells exhibits clear patterns in number and subtype organization. A single astrocyte precursor cell generates mostly ∼2-3 astrocytes and a single oligodendrocyte precursor cell generates largely ∼5-6 oligodendrocytes. Moreover, the output of individual precursor cells is largely of the same subtype and forms a local subcluster, suggesting that individual precursor cells are already restricted to a glial sublineage. Notably, the overall behavior of RGPs during gliogenesis in producing fate-restricted intermediate precursor cells that subsequently give rise to a relatively defined number of local progeny with similar properties is highly similar to that during neurogenesis with the generation of intermediate progenitor cells (Englund et al., 2005; Haubensak et al., 2004; Miyata et al., 2004; Sultan et al., 2018), indicating a common lineage design of RGPs in neurogenesis and gliogenesis.

Several studies have examined neocortical astrocyte generation at the clonal level using various techniques, such as transgenic labeling (Magavi et al., 2012), transposase (e.g., Star Track) labeling (Garcia-Marques and Lopez-Mascaraque, 2013; Siddiqi et al., 2014), and multi-addressable genome-integrative color (MAGIC) labeling (Clavreul et al., 2019). We have systematically examined the generation of astrocytes as well as oligodendrocytes by individual RGPs. Importantly, MADM labeling allows explicit assessment of the division behavior of labeled progenitors in vivo, thereby permitting direct capture of RGPs undergoing the neurogenesis-to-gliogenesis transition. While gliogenesis is generally thought to occur after neurogenesis, the exact timing of the transition remains uncertain. We found that the gliogenesis transition occur at different time points for individual RGPs, with a peak at ∼E16. Remarkably, regardless of the exact timing of transition, RGPs proceed to produce astrocytes, oligodendrocytes, or both in the consistent proportion of ∼60%:15%:25%, respectively, leading to the existence of three types of gliogenic clones. These results strongly suggest gliogenesis is an intrinsic and well-defined feature of RGPs.

The origin of astrocyte and oligodendrocyte lineages has been extensively debated (Richardson et al., 2006; Rowitch and Kriegstein, 2010). We found that ∼25% of gliogenic RGPs give rise to both astrocytes and oligodendrocytes, confirming the existence of bipotent oligodendrocyte-astrocyte progenitors in vivo. On the other hand, we did not find evidence of a distinct bipotent progenitor subtype in the neonatal neocortex, at least using the *Nestin-CreERT2* line. Instead, we observed fate-restricted astrocyte or oligodendrocyte precursor cells that generate a seemingly defined distribution of astrocytes or oligodendrocytes of the same subtype (L1A vs. PA vs. FA) or differentiation state (OPC vs. IMO vs. MO), as previously observed for rat optic nerve precursor cells in culture (Temple and Raff, 1986). Moreover, the progeny of individual astrocyte or oligodendrocyte precursor cells form spatially localized clusters, which likely contribute to the local glial cell heterogeneity (Bayraktar et al., 2020).

Notably, the number of astrocytes and/or oligodendrocytes in individual RGP clones exhibit substantial variability, as suggested in a recent study of astrocyte generation (Clavreul et al., 2019). We found that the variability largely stems from the number of intermediate astrocyte or oligodendrocyte precursor cells that individual RGPs generate. On average, individual gliogenic RGPs give rise to ∼2 astrocyte precursor cells and/or ∼2 oligodendrocyte precursor cells; yet, the exact number of intermediate precursor cells that individual RGPs generate appears to be stochastic. Such variability is likely critical for glial cell development and organization in the neocortex. Despite this variability, the glial output by individual RGPs exhibit obvious patterns in properties (i.e., astrocyte subtype or oligodendrocyte differentiation state). For example, we observed that individual RGPs generate L1A and PA or PA and FA, but not L1A and FA. Taken together, these results suggest that gliogenesis in the neocortex is organized.

While a substantial fraction (∼25%) of gliogenic RGPs generate both astrocytes and oligodendrocytes, their production at the single RGP level appears to be independent of each other. The number of astrocytes in RGP clones with both astrocytes and oligodendrocytes is the same as that in RGP clones with astrocytes only. Similarly, the number of oligodendrocytes in RGP clones with both oligodendrocytes and astrocytes is the same as that in RGP clones with oligodendrocytes only. These results suggest that astrogenesis and oligogenesis likely occur in a programmed manner. As long as an RGP generates astrocytes, it would execute the same astrogenesis program regardless of whether it also generates oligodendrocytes; and vice versa. As a consequence, RGPs generating both astrocytes and oligodendrocytes would undergo more cell divisions than those generating astrocytes or oligodendrocytes only. Notably, the number of neurons in glia-containing RGP clones is similar to that in non-glia-containing RGP clones, indicating the independence of neurogenesis and gliogenesis.

At both the single cell and population levels, we observed three types of gliogenic RGP clone in a consistent and reliable manner. It is unclear whether gliogenic RGPs are relatively homogenous (i.e., equipotent) with defined probabilities in generating astrocytes and/or oligodendrocytes or there are distinct subgroups of gliogenic RGPs destined to produce astrocytes, oligodendrocytes, or both. The defined proportion of RGPs with three different gliogenic transitioning modes is likely crucial for the generation of a proper number of astrocytes and oligodendrocytes in the neocortex. Similarly, it is unclear whether gliogenic and non-gliogenic RGPs are distinct subgroups. Taking advantage of the advances in single-cell profiling technologies, future studies will be required to explore the basis of diversity of gliogenic RGPs and the regulation of neocortical gliogenesis.

Oligodendrocytes in the neocortex have multiple developmental origins and are generated by RGPs in the developing neocortex as well as ventral telencephalon (Kessaris et al., 2006). We noticed that oligodendrocytes labeled in this study (i.e., neocortical RGP-derived) are more abundantly located in the deep layers and white matter, raising the possibility that the ventral telencephalon RGP-derived oligodendrocytes are more frequently distributed in the superficial layers. Recent comprehensive single cell analysis revealed that glial cells in different layers exhibit distinct transcriptional profiles (Bayraktar et al., 2020; Lanjakornsiripan et al., 2018; Marques et al., 2016). This diversity of glial cells may be rooted partly in their developmental origins. In line with this, individual astrocyte or oligodendrocyte precursor cells produce astrocytes or oligodendrocytes with similar properties, forming local clusters. Together, these results suggest that progenitor and lineage origin influence the number, subtype and spatial distribution of glial cells in the neocortex, in conjunction with local environment.

While glial cells and neurons share the same progenitor cell origin of RGPs, removal of NF1, a tumor suppressor (Ratner and Miller, 2015), in RGPs does not affect neurogenesis, but disrupts gliogenesis. This is consistent with the fact that, at the single RGP level, gliogenesis and neurogenesis occur in separate developmental time windows. Moreover, the molecular control of gliogenesis and neurogenesis can be distinct. Furthermore, while astrocyte and oligodendrocyte generation are both enhanced upon NF1 removal, the extent of change is clearly different. The average number of astrocytes generated by a single gliogenic RGP increases by ∼2-fold in the absence of NF1. In comparison, the average number of oligodendrocyte increases by >15-fold. Strikingly, we observed individual *Nf1* mutant clones with hundreds to thousands of immature, proliferative OPCs. On the other hand, we did not observe any extraordinarily large *Nf1* mutant clones of astrocytes, despite that far more astrocyte clones were labeled than oligodendrocyte clones. Interestingly, NF1 loss leads to distinct changes in astrogenesis and oligogenesis by individual RGPs. For astrogenesis, NF1 loss causes an increase in astrocyte precursor cell generation, but not the output of individual astrocyte precursor cells. On the other hand, for oligogenesis, NF1 loss results in an increase in OPC generation as well as the output of individual OPCs. The increase in individual OPC output is especially drastic. Given that *Nf1* mutation is linked to brain tumorigenesis, such as pediatric glioma (Anastasaki et al., 2017), these results suggest that oligodendrocyte lineage is particularly susceptible to tumor formation, as indicated in previous studies (Ligon et al., 2007; Liu et al., 2011). Notably, recent studies suggest that brain tumorigenesis often reflects the abnormal activation of the early developmental processes (Vladoiu et al., 2019). Further interrogation of the abnormal behavior of the oligodendrocyte lineage, especially OPCs, will greatly advance our understanding of the cellular origin and basis of brain tumor formation.

## ACKNOWLEDGEMENTS

We thank Dr. Nicolette Kessaris (University College London, UK), Dr. Ryoichiro Kageyama (Kyoto University, Japan), Dr. Simon Hippenmeyer (IST, Austria), and Dr. Yuan Zhu (The Children’s Research Institute, USA) for providing *Emx1-CreERT1*, *Nestin-CreERT2*, *MADM11*, and *Nf1^fl/fl^* mice, respectively; We thank Shi lab members and Andy Yu Shi for their input and comments on the manuscript. This work was supported by grants from Beijing Outstanding Young Scientist Program (BJJWZYJH01201910003012 to S.-H.S.), NIH (R01DA024681 to S.-H.S.) and the Howard Hughes Medical Institute (to S.-H.S.). B.D.S acknowledges funding from the Royal Society E.P. Abraham Research Professorship (RP\R1\180165) and Wellcome Trust (098357/Z/12/Z).

## AUTHOR CONTRIBUTIONS

Z.S., Y.L. and S.-H.S. conceived, and B.D.S. and S.-H.S. supervised the project; Z.S. and Y.L. performed most of the experiments with help from Y.P., X.Z., J. M. and L.H.; J.Y., D.J.J. and Y.X. performed quantitative analysis of the datasets; and Z.S. and S.-H.S. wrote the manuscript with inputs from all authors.

## DATA AVAILABILITY

The datasets generated during and/or analysed during the current study are available from the corresponding author on reasonable request. The authors have no competing financial or non-financial interests.

## EXPERIMENTAL PROCEDURE

### Mouse Lines

*MADM-11GT* (stock# 013749) and *MADM-11TG* (stock# 013751) mouse lines (Hippenmeyer et al., 2010) were obtained from Jackson Laboratory (Bar Harbor, ME); *Emx1-CreERT2* (Kessaris et al., 2006), *Nestin-CreERT2* (Imayoshi et al., 2006), and *Nf1^fl/fl^* (Zhu et al., 2001) mouse lines were kindly provided by Dr. Nicoletta Kessaris, Dr. Ryoichiro Kageyama, and Dr. Yuan Zhu, respectively. CD-1 mice were obtained from Beijing Vital River Laboratory Animal Technology Co., Ltd (Beijing, China). *Nf1* floxed allele was genetically linked to *MADM-11GT* through meiotic recombination. Genotyping was carried out using standard PCR protocols. Both male and female mice were used in the study. For clone induction at the embryonic stage, pregnant females were either injected intraperitoneally (E13 or E14; 10-20 μg/g of body weight) or orally gavaged (E15, E16 or E17; 200∼400 μg/g of body weight) with Tamoxifen (T5648, Sigma) dissolved in corn oil (C8267, Sigma). Live embryos were recovered at E18–E19 through cesarean section, fostered, and raised for further analysis. For clone induction at the neonatal stage, pups were injected intraperitoneally with Tamoxifen dissolved in corn oil at a dose of 100-200 μg/g of body weight. The mice were maintained at the facility of Tsinghua University, and all animal procedures were approved by the Institutional Animal Care and Use Committee (IACUC). For timed pregnancies, the plug date was designated as E0 and the date of birth was defined as P0. No wild animal or field-collected sample was used in the study.

### Brain Sectioning, Immunohistochemistry, Imaging, and 3D Reconstruction

Mice were perfused intracardially with ice-cold phosphate-buffered saline (PBS, pH 7.4), followed by 4% paraformaldehyde (PFA) in PBS (pH 7.4). Brains were post-fixed with 4% PFA at 4 °C for ∼6 hours, cryo-protected, and sectioned at 60 or 100 μm using microtome (Leica Microsystems) for immunohistochemistry, as previously described (Gao et al., 2014)). The following primary antibodies were used: chicken anti-GFP (GFP-1020; 1:1000; Aves), rabbit anti-RFP (600-401-379; 1:500; Rockland), rabbit anti-GFAP (G9269; 1:100; Sigma), rat anti-NG2 (546930; 1:100; ThermoFisher), mouse anti-OLIG2 (MABN50; 1:500; Millipore), rabbit anti-OLIG2 (AB9610; 1:500; Millipore), mouse anti-S100β (MA1-25005; 1:100; ThermoFisher), mouse anti-MBP (ab62631; 1:500; Abcam) and mouse anti-NEUN (MAB377; 1:500; Millipore). Alexa flour 488-Donkey anti-chicken (703-546-155; 1:1000, Jackson ImmunoResearch), Alexa flour 488-Donkey anti-rat (A21208; 1:1000, ThermoFisher), Alexa flour 555-Donkey anti-rabbit (A31572; 1:1000, ThermoFisher), Alexa flour 647-Donkey anti-mouse (A31571; 1:1000, ThermoFisher), Alexa flour 647-Donkey anti-rabbit (A31573; 1:1000, ThermoFisher) secondary antibodies were used.

Brain sections were mounted on glass slides, imaged using confocal microscopy (FV3000, Olympus) or slide scanner (Axio Z1, Zeiss), and reconstructed using Neurolucida (MBF Bioscience). For 3D reconstruction, each section was analyzed sequentially in the rostral to caudal order. The boundaries of the entire brain and lateral ventricles were traced and aligned. Individual labeled neurons, astrocytes, and oligodendrocytes were represented as colored symbols (three to four times the size of the cell body). Layer boundaries based on nuclear staining were also documented. Cortical areas were identified based on the Allen Brain Atlas (http://mouse.brain-map.org/static/atlas). Images were analyzed by ZEN (Zeiss), FV31S-DT (Olympus), IMARIS (Bitplane), or Photoshop (Adobe).

### NND Analysis and Cluster/Subcluster Identification

The distribution of the nearest neighbor distance (NND) reflects the spatial point pattern of the data set, as previously described (Diggle, 2003). Specifically, given N cells in a data set, for each cell *i* the distance to its closest neighbor was measured and denoted as ***d_i_***, the NND for cell ***i***. The indicator function *f(y,d)* was then calculated as:

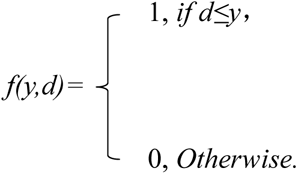

Thus, the cumulative distribution function (CDF) of NND is:

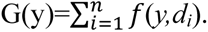

In this analysis, the shorter NND distance reflects clustering, whereas the longer NND distance reflects dispersing. Simulated random data sets contained the same number of data points within the same volume (i.e., the neocortex) as the experimental data sets, and were repeated 100 times.

Cluster identification in neonatal clones and subcluster identification in embryonic clones were performed using mean shift analysis, a clustering method without the requirement of specifying the exact cluster number, thereby permitting to analyze all experimental data with consistent spatial parameters. The process started from a random data point/cell within the dataset. Vectors from the starter cell to every other cell with a smaller distance than the detection bandwidth were calculated. The cluster center then shifted along the vector and the calculation repeated till the cluster center reached a stable density center. Afterwards, a new center randomly selected from the dataset not reached by the previous cluster center was used to repeat the next analysis loop till every cell in the dataset was clustered into clusters. Finally, the cluster centers closer than the merge threshold were merged together. The spatial parameters of the mean shift analysis were determined using neonatal glial clones. NND analysis and subcluster analysis were measured using Matlab (R2016b, MathWorks).

### Quantification and Statistical Analysis

Mice subjected to the analyses were littermates, age-matched, and includes both sexes. Sample size was determined to be adequate based on the magnitude and consistency of measurable differences between groups. Statistical significance was determined using Chi-square or two-sided non-parametric Mann-Whitney-Wilcoxon test or Kolmogorov-Smirnow test, and the test results were given as exact values in the figures. Statistical significance was defined as *p* < 0.05. Statistical tests were performed with Prism (version 7, GraphPad). Effect sizes were calculated using Pearson’s r (Chi-square) or U/(n1*n2) (Mann-Whitney-Wilcoxon test). Values in bar graphs indicate mean ± SEM.

## SUPPLEMENTAL FIGURE LEGENDS

**Figure S1.**
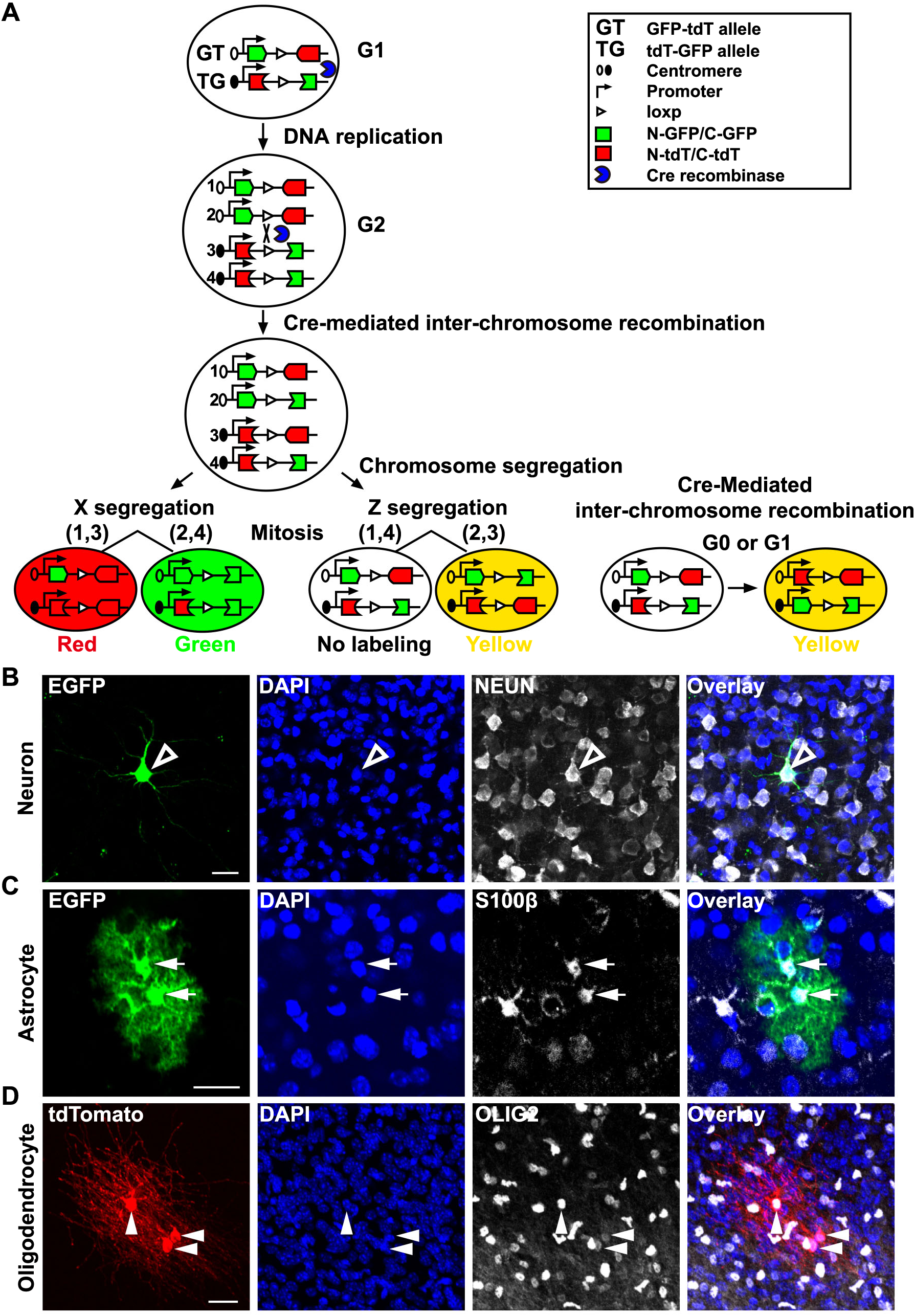
**MADM-based clonal analysis of neocortical gliogenesis by RGPs. Related to Figure 1.** (A) Schematic of MADM labeling. Cre recombinase-mediated interchromosomal recombination in the G_2_ phase of dividing progenitor followed by X-segregation (G_2_-X, segregation of recombinant sister chromatids into separate daughter cells) reconstitutes one of two fluorescent markers, EGFP (green) or tdTomato (red), in each of the two daughter cells. As such, G_2_-X MADM events result in permanent and distinct labeling of the two descendent lineages, thereby allowing a direct assessment of the division pattern (symmetric vs. asymmetric) and potential (the number of progeny) of the original dividing progenitor. In addition, upon G_2_-Z (congregation of recombinant sister chromatids into the same daughter cell), G_1_, or G_0_ recombination events, green and red (i.e., yellow) fluorescent proteins are restored simultaneously in the same cell, resulting in double-labeled (yellow) cells. (B) Confocal images of a representative MADM-labeled excitatory neuron (open arrowhead) stained with antibodies against EGFP (green) and NEUN (white), and with DAPI (blue). Scale bar: 20 μm. (C) Confocal images of representative MADM-labeled astrocytes (arrows) stained with antibodies against EGFP (green) and S100β (white), and with DAPI (blue). Scale bar: 20 μm. (D) Confocal images of representative MADM-labeled oligodendrocytes (arrowheads) stained with antibodies against tdTomato (red) and OLIG2 (white), and with DAPI (blue). Scale bar: 20 μm.

**Figure S2.**
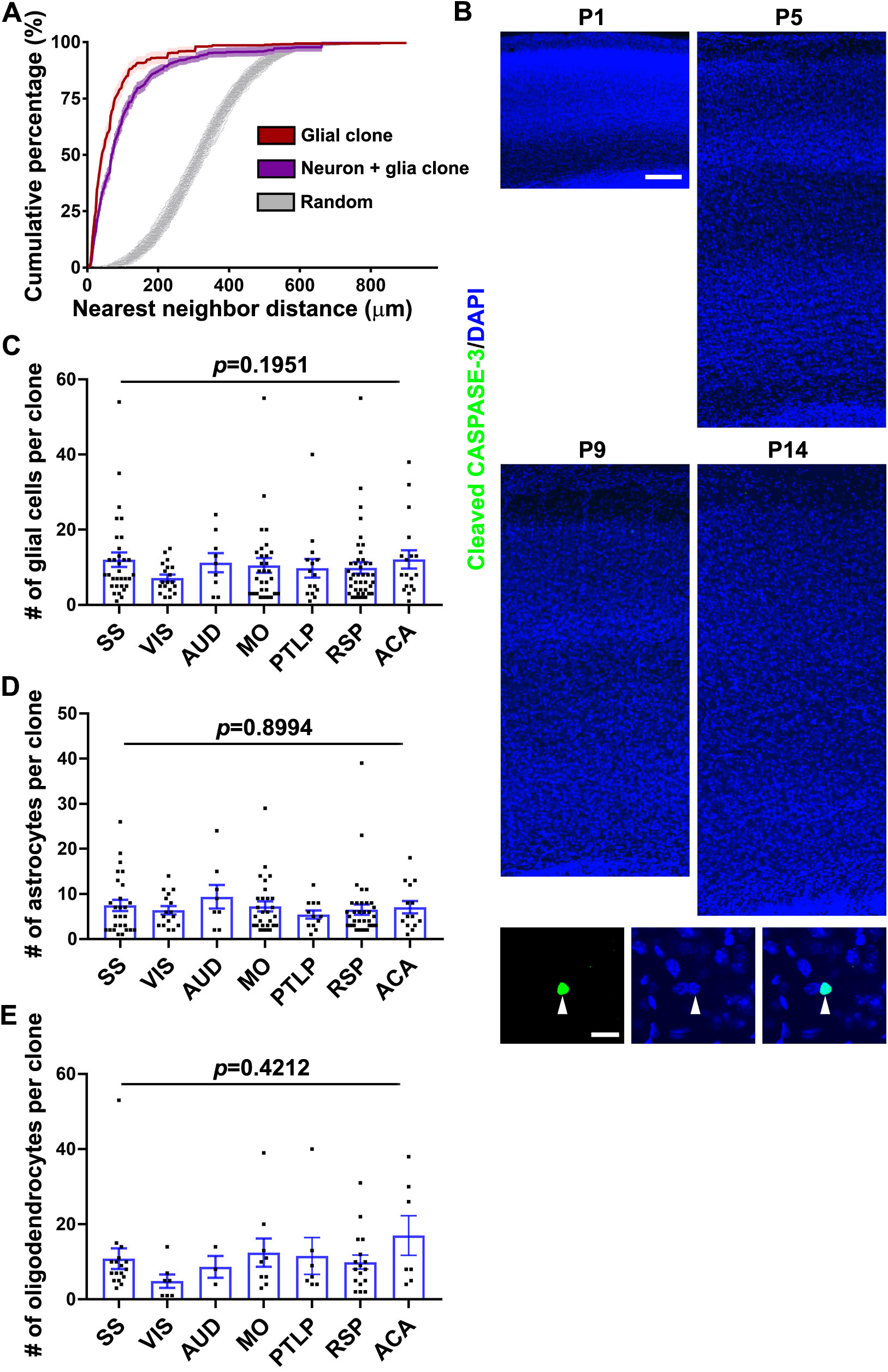
**Clustering and consistent size of glial cell clone. Related to Figure 1.** (A) NND analysis of MADM-labeled clonally related glial cells (red) and glial cells and neurons (magenta) labeled at E13 and analyzed at P21-34 in comparison with randomly simulated control datasets with the same number of data points in the same volume (grey). (B) Representative confocal images of neocortical sections stained with Cleaved CASPASE-3 (green) and with DAPI (blue) at different postnatal stages. Zoom in images of a Cleaved CASPASE-3-expressing cell are shown at the bottom. Scale bars: 100 μm and 20 μm. (C) Quantification of the number of glial cell in individual G_2_-X N + G clones located in different neocortical areas (SS, n = 32; VIS, n = 19; AUD, n = 9; MO, n = 30; PTLP, n = 15; RSP, n = 41; ACA, n = 18). SS, somatosensory cortex; VIS, visual cortex; AUD, auditory cortex; MO, motor cortex; PTLP, posterior parietal association areas; RSP, retrosplenial area; ACA, anterior cingulate area. Bar plots and lines indicate mean ± SEM and dots indicate individual clones. Statistical analysis was performed using two-sided Mann-Whitney-Wilcoxon test. (D) Quantification of the number of astrocyte in individual G_2_-X N + G clones with astrocyte located in different neocortical areas (SS, n = 27; VIS, n = 16; AUD, n = 8; MO, n = 28; PTLP, n = 12; RSP, n = 36; ACA, n = 14). Bar plots and lines indicate mean ± SEM and dots indicate individual clones. Statistical analysis was performed using two-sided Mann-Whitney-Wilcoxon test. (E) Quantification of the number of oligodendrocyte in individual G_2_-X N + G clones with oligodendrocyte located in different neocortical areas (SS, n = 17; VIS, n = 7; AUD, n = 3; MO, n = 9; PTLP, n = 7; RSP, n = 17; ACA, n = 7). Bar plots and lines indicate mean ± SEM and dots indicate individual clones. Statistical analysis was performed using two-sided Mann-Whitney-Wilcoxon test.

**Figure S3.**
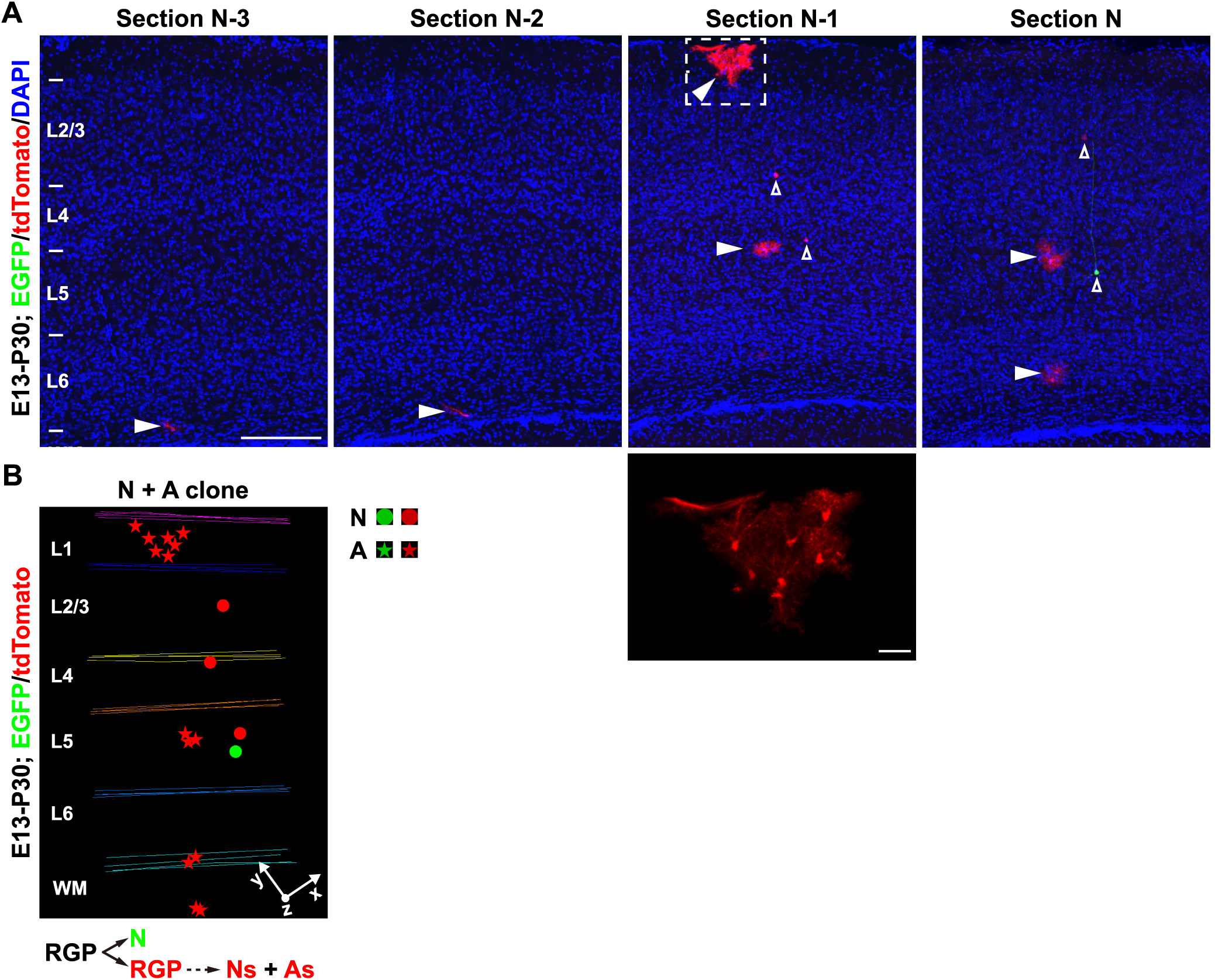
**MADM labeling of RGP clone with neuron and astrocyte (N + A). Related to Figure 2-4.** (A) Confocal images of a representative E13-P30 G_2_-X N + A clone. Arrowheads indicate astrocytes and open arrowheads indicate excitatory neurons. High-magnification image of labeled astrocytes (broken lines) is shown in inset. Scale bars: 200 μm and 40 μm. (B) 3D reconstruction image of the clone shown in A. The division pattern and progeny output are shown at the bottom. Layers are shown to the left. Colored lines indicate the brain contours and layer boundaries. The x/y/z axes indicate the spatial orientation of the clone with the y axis parallel to the brain midline and pointing dorsally.

**Figure S4.**
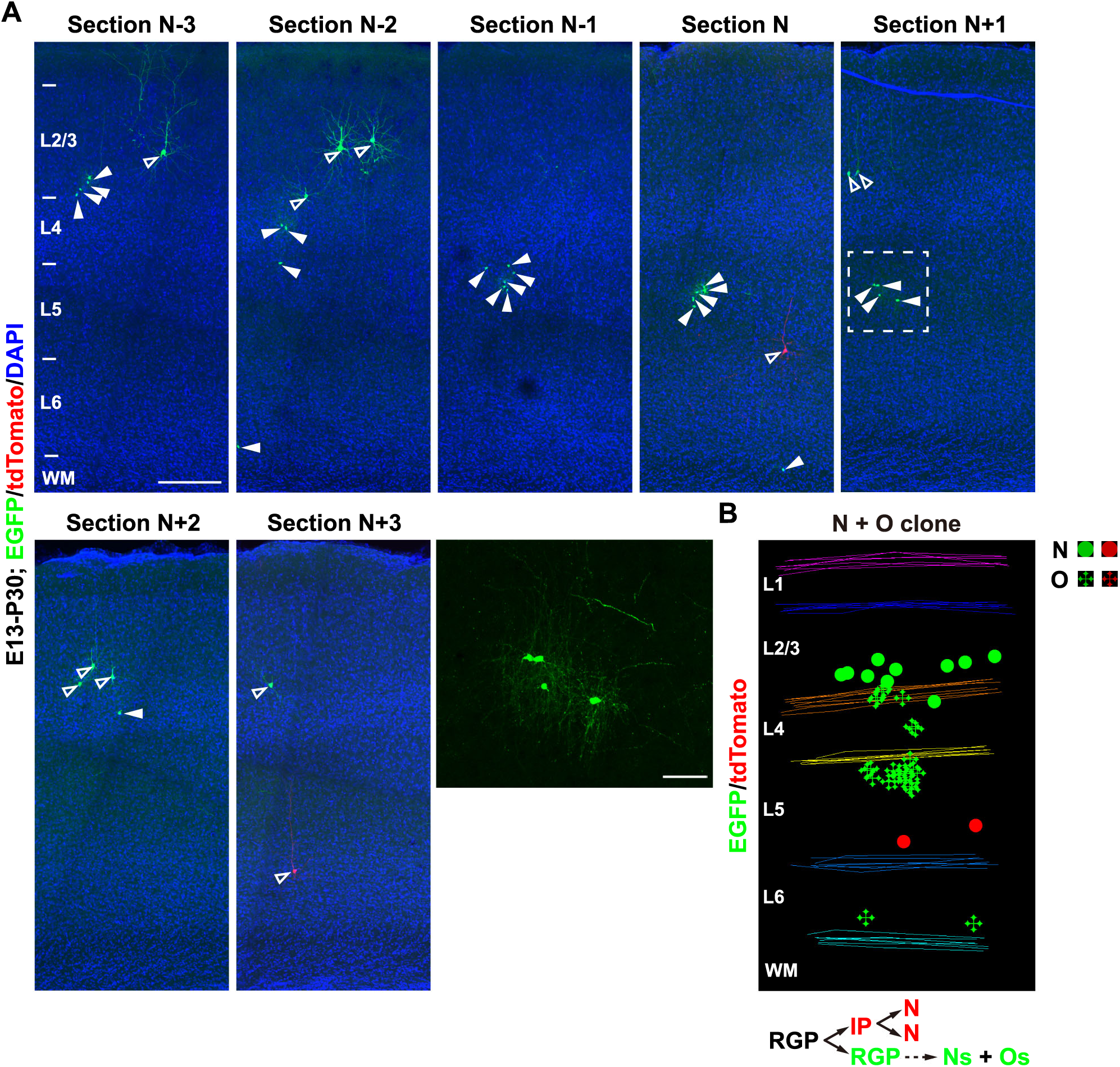
**MADM labeling of RGP clone with neuron and oligodendrocyte (N + O). Related to Figure 2-4.** (A) Confocal images of a representative E13-P30 G_2_-X N + O clone. Arrowheads indicate oligodendrocytes and open arrowheads indicate excitatory neurons. High-magnification image of labeled oligodendrocyte (broken lines) is shown in inset. Scale bars: 200 μm and 20 μm. (B) 3D reconstruction image of the clone shown in A. The division pattern and progeny output are shown at the bottom.

**Figure S5.**
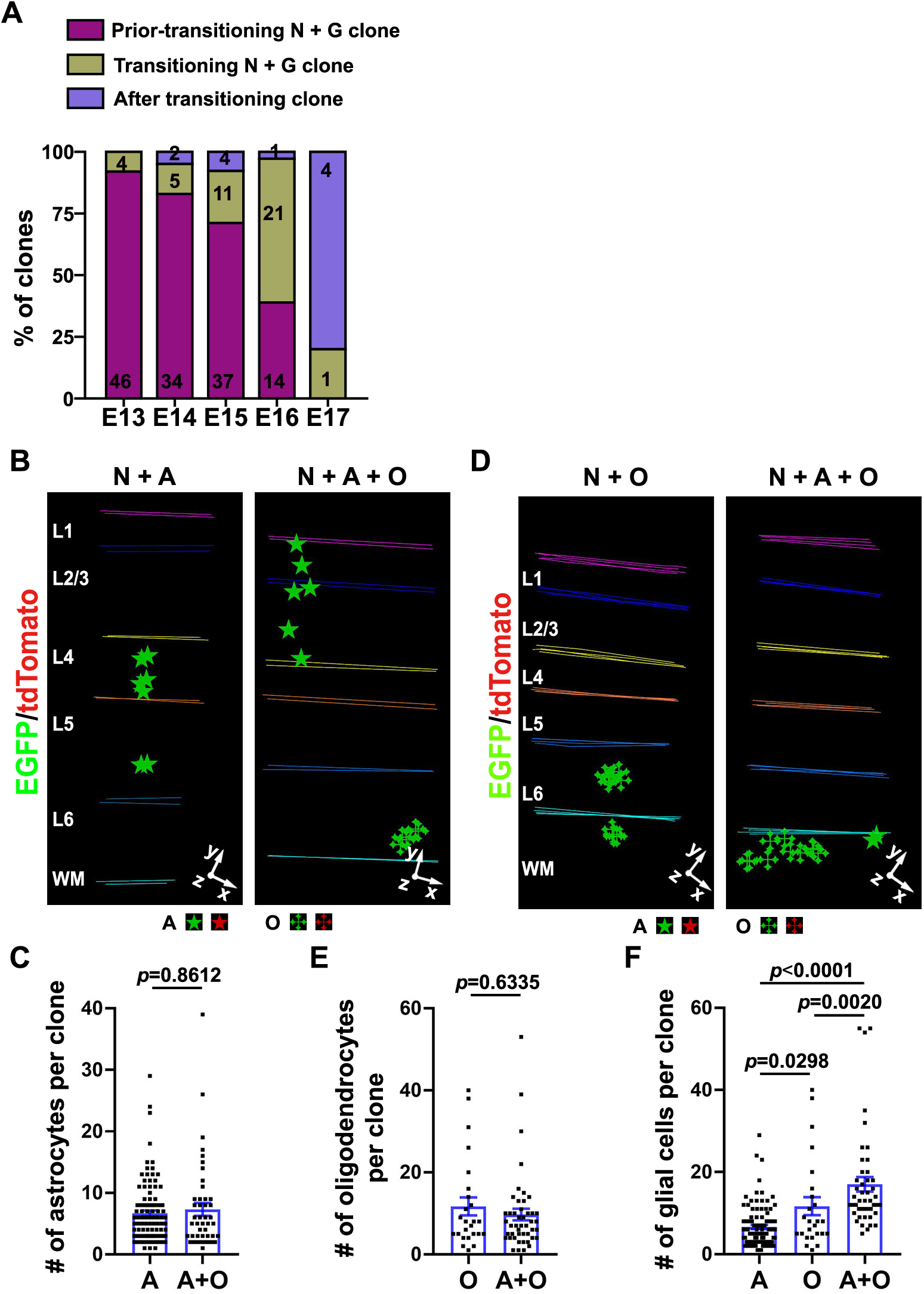
**Independent generation of astrocyte and oligodendrocyte by individual RGPs. Related to Figure 3.** (A) Quantification of the fraction of non-transitioning N + G clone, transitioning N + G clone, and glia only clone labeled at E13-E17. (B) 3D reconstruction images of representative G_2_-X N + A and N + A + O clones. Neurons are not shown in the images. (C) Quantification of the number of astrocyte in N + A (A, n = 101) and N + A + O (A + O, n = 46) clones. Bar plots and lines indicate mean ± SEM and dots indicate individual clones. Statistical analysis was performed using two-sided Mann-Whitney-Wilcoxon test. Note that the numbers of astrocyte in A and A + O clones are similar. (D) 3D reconstruction images of representative G_2_-X N + O and N + A + O clones. Neurons are not shown in the images. (E) Quantification of the number of oligodendrocyte in N + O (O, n = 25) and N + A + O (A + O, n = 46) clones. Bar plots and lines indicate mean ± SEM and dots indicate individual clones. Statistical analysis was performed using two-sided Mann-Whitney-Wilcoxon test. Note that the numbers of oligodendrocyte in O and A + O clones are similar. (F) Quantification of the number of glial cell in A (n = 101), O (n = 25), and A + O (n = 46) clones. Bar plots and lines indicate mean ± SEM and dots indicate individual clones. Statistical analysis was performed using two-sided Mann-Whitney-Wilcoxon test. Note that the number of glial cell in the A + O clone is approximately the sum of those in the A and O clones.

**Figure S6.**
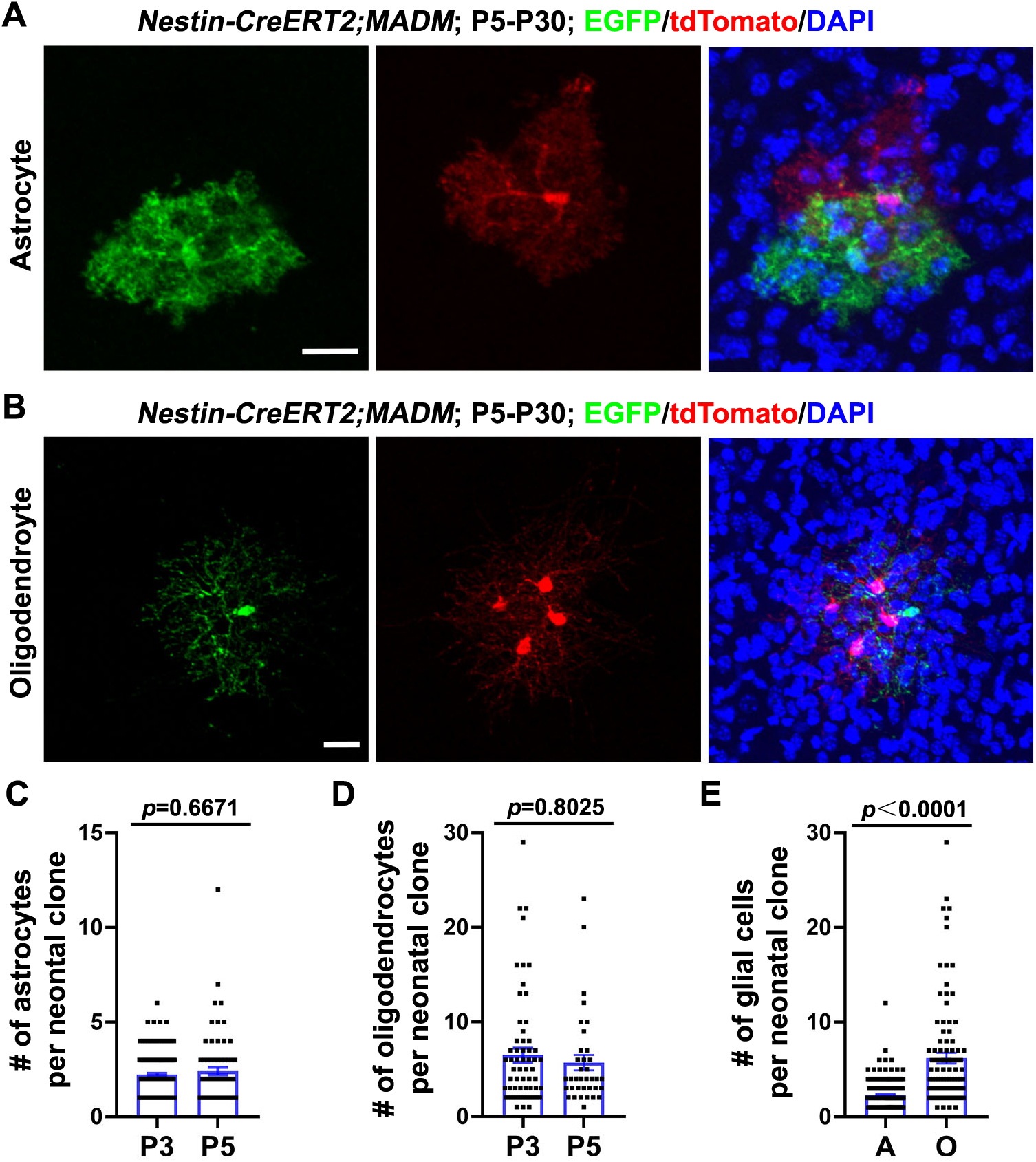
**MADM labeling of glial clones originated from fate-restricted precursor cells labeled at the neonatal stage. Related to Figure 4.** (A) Confocal images of a representative astrocyte only G_2_-X clone labeled at P5 and examined at P30. Brain sections were stained with antibodies against EGFP (green) and tdTomato (red), and with DAPI (blue). Scale bar: 20 μm. (B) Confocal images of a representative oligodendrocyte only G_2_-X clone labeled at P5 and examined at P30. Brain sections were stained with antibodies against EGFP (green) and tdTomato (red), and with DAPI (blue). Scale bar: 20 μm. (C) Quantification of the number of astrocyte in individual neonatal G_2_-X clones with astrocyte labeled at P3 (n = 197) and P5 (n = 76). Bar plots and lines indicate mean ± SEM and dots indicate individual clones. Statistical analysis was performed using two-sided Mann-Whitney-Wilcoxon test. (D) Quantification of the number of oligodendrocyte in neonatal G_2_-X clones with oligodendrocyte labeled at P3 (n = 71) and P5 (n = 36). Bar plots and lines indicate mean ± SEM and dots indicate individual clones. Statistical analysis was performed using two-sided Mann-Whitney-Wilcoxon test. (E) Quantification of the number of astrocyte or oligodendrocyte in neonatal G_2_-X clones (astrocyte, n = 273; oligodendrocyte, n = 107). Bar plots and lines indicate mean ± SEM and dots indicate individual clones. Statistical analysis was performed using two-sided Mann-Whitney-Wilcoxon test.

**Figure S7.**
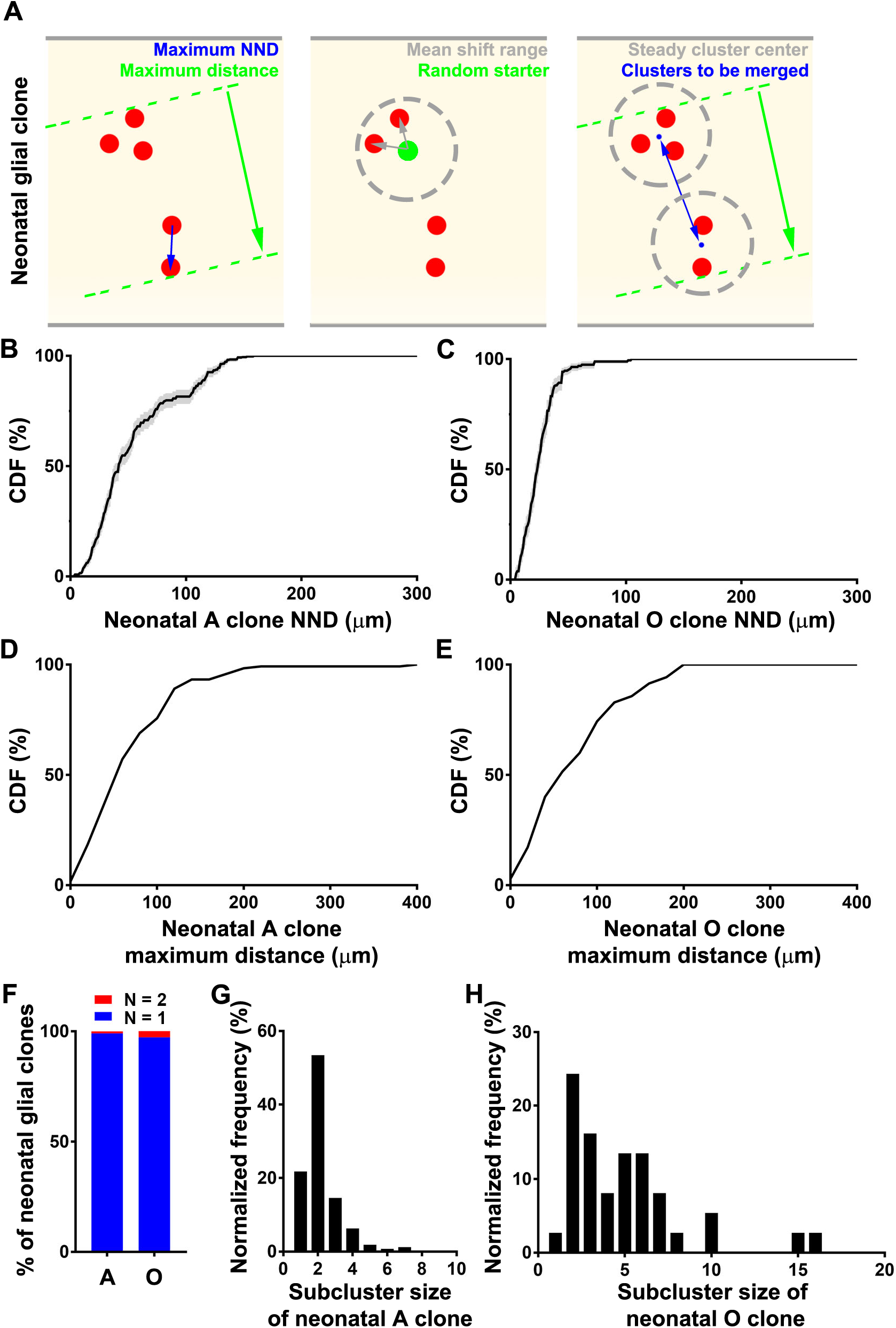
**Mean shift analysis for identifying glial subclusters in individual RGP clones. Related to Figure 4.** (A) Diagram of mean shift clustering method and parameter determination in analyzing glial neonatal clones. Left: Maximum NND of neonatal glial clones is used as mean shift detection range bandwidth to search for neighbor cells from a random starter. Middle: Calculate the average of vectors from the center to every cell within the detection region and shift the center to make the mean vector minimum. Right: Merge the clusters to cover the entire neonatal clone. The threshold of the cluster center distance to be merged should be no more than the difference between the maximum clone distance and the mean shift bandwidth. (B) Cumulative distribution function (CDF) of the NND of neonatal astrocyte clones. The onset of the plateau value was used as the distance threshold of the cluster centers to be merged. (C) CDF of the NND of neonatal oligodendrocyte clones. The onset of the plateau value was used as the distance threshold of the cluster centers to be merged. (D) CDF of the maximum distance of neonatal astrocyte clones. The onset of the plateau value was used as the mean shift detection bandwidth. (E) CDF of the maximum distance of neonatal oligodendrocyte clones. The onset of the plateau value was used as the mean shift detection bandwidth. (F) Quantification of the subcluster number of astrocyte (A) or oligodendrocyte (O) in individual neonatal glial clones identified using the mean shift clustering method. Notice that the vast majority of clones contained only one cluster. (G) Histogram of the subcluster size of neonatal astrocyte clones. (H) Histogram of the subcluster size of neonatal oligodendrocyte clones.

**Figure S8.**
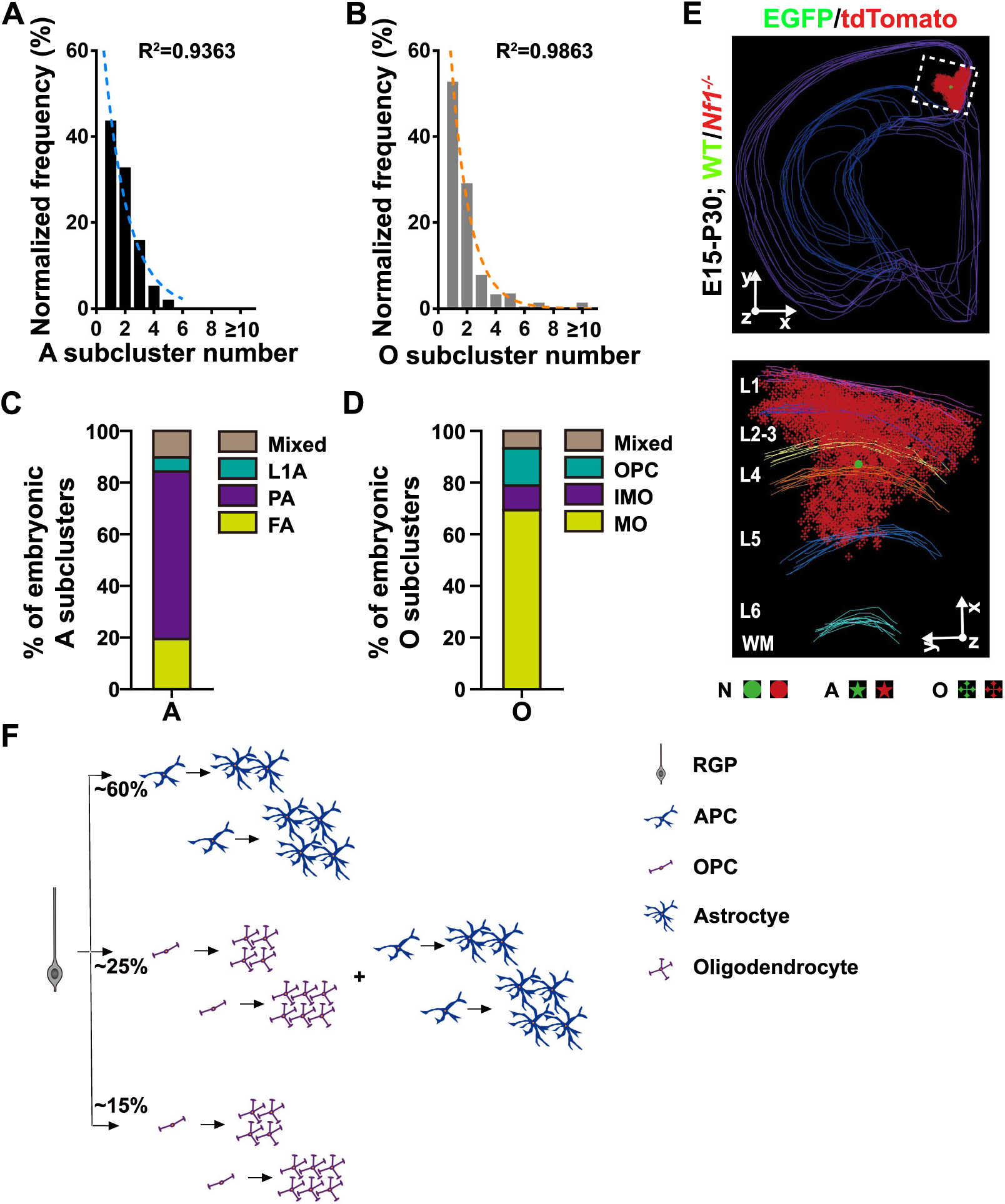
**Glial cell properties in individual subclusters of RGP clone, clones with an extraordinarily large number of *Nf1* mutant OPCs and neocortical gliogenesis program by RGPs. Related to Figure 4 and 7.** (A) Histogram of the number of astrocyte subcluster in individual embryonic clones (n = 147). Note that the distribution can be well approximated by a geometric distribution (blue broken line), indicating that the generation of astrocyte precursor cell by individual RGPs is stochastic. (B) Histogram of the number of oligodendrocyte subcluster in individual embryonic clones (n = 71). Note that the distribution can be well approximated by a geometric distribution (yellow broken line), indicating that the generation of oligodendrocyte precursor cell by individual RGPs is stochastic. (C) Quantification of the astrocyte subtype composition of individual subclusters identified in the embryonic G_2_-X N + G clone with astrocyte (n = 271). Note that the vast majority of subclusters contain the same subtype of astrocyte. (D) Quantification of the oligodendrocyte differentiation state of individual subclusters identified in the embryonic G_2_-X N + G clone with oligodendrocyte (n = 121). Note that the vast majority of subclusters contain the same differentiation state of oligodendrocyte. (E) 3D reconstruction image of a representative clone with excessive number of *Nf1^-/-^* OPCs. (F) Schematic summary of neocortical gliogenesis program by RGPs.

## SUPPLEMENTAL MOVIE LEGEND

**Movie S1. 3D reconstruction of a brain hemisphere with only one G2-X N + G clone labeled in the neocortex; Related to Figure 1.** Colored lines indicated the brain contours, colored dots represent the cell bodies of labeled excitatory neurons, colored stars indicate the cell bodies of labeled astrocytes, and colored crosses indicate the cell bodies of labeled oligodendrocytes.

